# Biological mechanisms of aging predict age-related disease multimorbidities in patients

**DOI:** 10.1101/2021.05.04.442567

**Authors:** Helen C Fraser, Valerie Kuan, Ronja Johnen, Magdalena Zwierzyna, Aroon D Hingorani, Andreas Beyer, Linda Partridge

## Abstract

Genetic, environmental and pharmacological interventions into the aging process can confer resistance to a multiple age-related diseases in laboratory animals, including rhesus monkeys. These findings imply that mechanisms of aging might contribute to patterns of multimorbidity in humans, and hence could be targeted to prevent multiple conditions simultaneously. To address this question, we text mined 917,645 literature abstracts followed by manual curation, and found strong, non-random associations between age-related diseases and aging mechanisms, confirmed by gene set enrichment analysis of GWAS data. Integration of these associations with clinical data from 3.01 million patients showed that age-related diseases associated with each of five aging mechanisms were more likely than chance to be present together in patients. Genetic evidence revealed that innate and adaptive immunity, the intrinsic apoptotic signalling pathway and activity of the ERK1/2 pathway played a significant role across multiple aging mechanisms and multiple, diverse age-related diseases. Mechanisms of aging therefore contribute to multiple age-related diseases and to patterns of human age-related multimorbidity, and could potentially be targeted to prevent more than one age-related condition in the same patient.

## Introduction

Age-associated accumulation of molecular and cellular damage leads to an increased susceptibility to loss of function, disease and death^1^. Aging is the major risk factor for many chronic and fatal human diseases, including Alzheimer’s disease, multiple cancers, cardiovascular diseases and type 2 diabetes mellitus (T2DM), which are collectively known as age-related diseases (ARDs)^2^. However, genetic^3^, environmental^4^ and pharmacological^5^ interventions can ameliorate loss of function during aging and confer resistance to multiple age-related diseases in laboratory animals. Age-related multimorbidity, the presence of more than one ARD in an individual, is posing a major and increasing challenge to health care systems worldwide^6^. An important, open question, therefore, is whether mechanisms of aging in humans contribute to multimorbidity in patients, and hence whether intervention into these mechanisms could prevent or treat more than one ARD simultaneously^7^.

Specific biological mechanisms begin to fail as an individual ages^1^. Nine major aging processes were summarised as *“The Hallmarks of Aging”*^1^: genomic instability, telomere shortening, epigenetic changes, impaired protein homeostasis, impaired mitochondrial function, deregulated nutrient sensing, cellular senescence, exhaustion of stem cells and altered intercellular communication (Fig 1). Individual aging hallmarks are present in the development or disordered physiology of specific ARDs^8^. For example, loss of proteostasis appears to have a prominent role in neurodegenerative disorders, such as Alzheimer’s and Parkinson’s diseases, which are associated with protein aggregates composed of amyloid-beta and *α*-synuclein, respectively^9^. Genomic instability and epigenetic alterations frequently contribute to development of cancers of, for example, the breast and bowel^10^. The role of genes in individual human ARDs and ARD multimorbidity has been studied extensively^11–13^, as has the link between aging hallmarks and individual ARDs^11,14^. For example, previous studies have demonstrated that multiple, individual human ARDs share gene ontology (GO) terms linked to mechanisms of aging, specifically aging hallmarks^11^. However, whether these underlying mechanisms of aging contribute to the occurrence of multimorbidity in patients has not previously been investigated. Here, we explore the notion that the same aging hallmark may contribute to risk of multiple ARDs and, therefore, results in their co-occurrence in the same individual (i.e., ARD multimorbidity). In model organisms, altering the activity of signalling pathways, such as the insulin/ insulin-like growth factor signalling (IIS) pathway^1^, Ras-ERK pathway^15^, immune pathways^16^ or p53 pathways^17^, can delay ARD and/ or extend lifespan. These pathways are also intertwined with the aging hallmarks^1^. Therefore, we also explored the notion that common signalling pathways are shared across all aging hallmarks and, thus, multiple, aging hallmark-associated ARDs.

**Figure 1.**
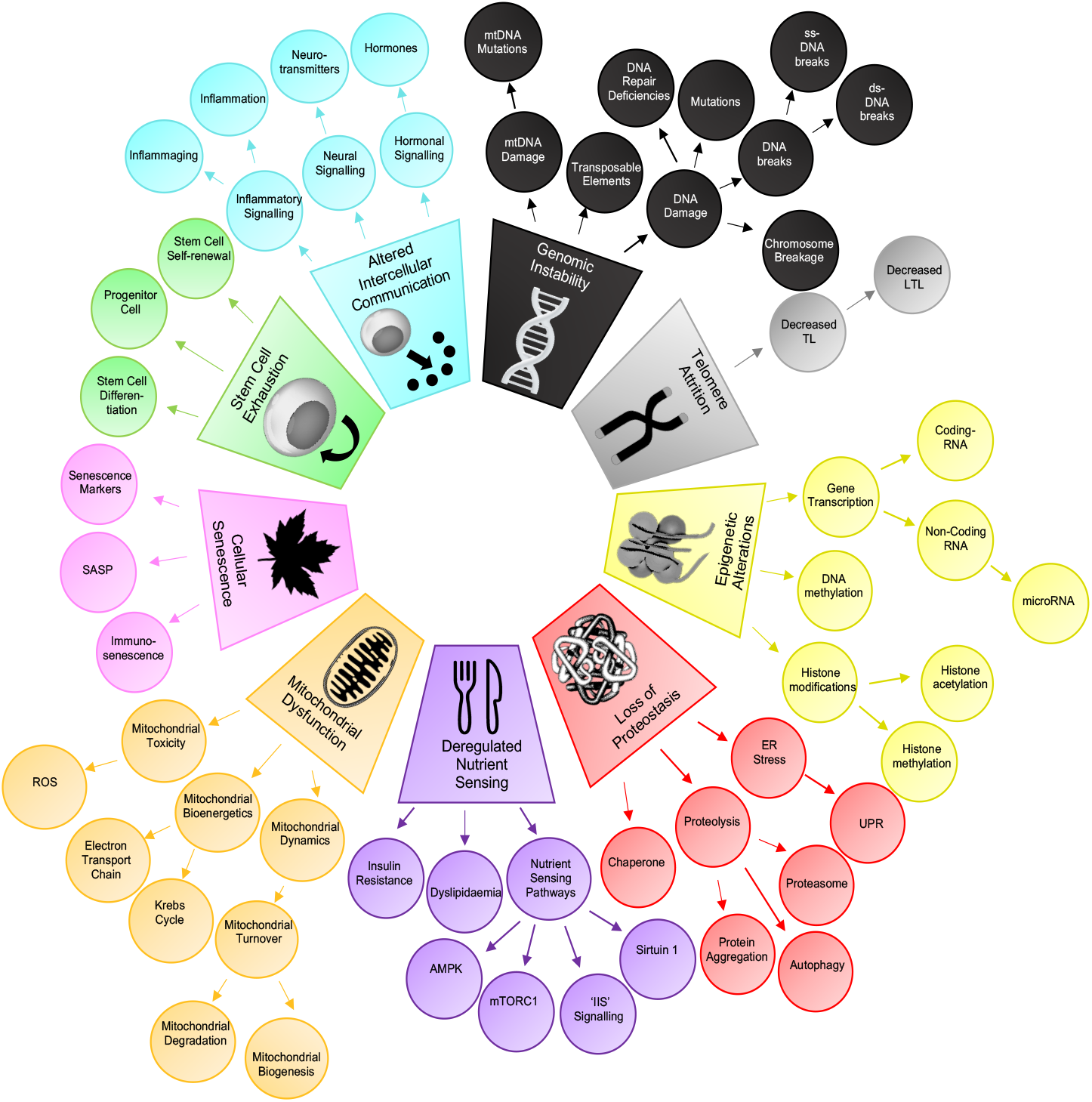
The “Hallmarks of Aging” expanded into a taxonomy. The nine original aging hallmarks were expanded into a taxonomy of 65 related terms and four levels. Figure adapted from López-Otin *et al*. (2013). Abbreviations: Table S9.

We integrated evidence derived from the scientific literature abstracts, genome wide association (GWA) studies and electronic health records to explore the role of aging hallmarks in human ARD multimorbidities. We began by scoring co-mentions of aging hallmarks and ARDs in 917,645 scientific literature abstracts and verified the aging hallmark-ARD associations that emerged using manual curation. Using the scores of verified literature aging hallmark-ARD associations, we identified the top 30 ranked ARDs (i.e., top 30 ARDs) associated with each aging hallmark (Fig 2a). To confirm these associations independently, we used publicly available GWAS data to explore whether annotations of proteins encoded by genes associated with the top 30 ARDs were enriched for processes related to the same hallmark (Fig 2b). GO annotation of the GWAS data also indicated that diverse, aging hallmark-associated ARDs were linked with common signalling pathways. Next, clinical data from 3.01 million patients was used to construct networks of aging-related multimorbidities, which were developed previously^18,19^. We used the patient data to examine whether the top 30 ARDs associated with each aging hallmark were more frequent multimorbidities in individual patients than expected by chance (Fig 2c). We also investigated whether aging hallmarks contribute to ARDs with incompletely understood mechanisms of development (Fig 2d).

**Figure 2.**
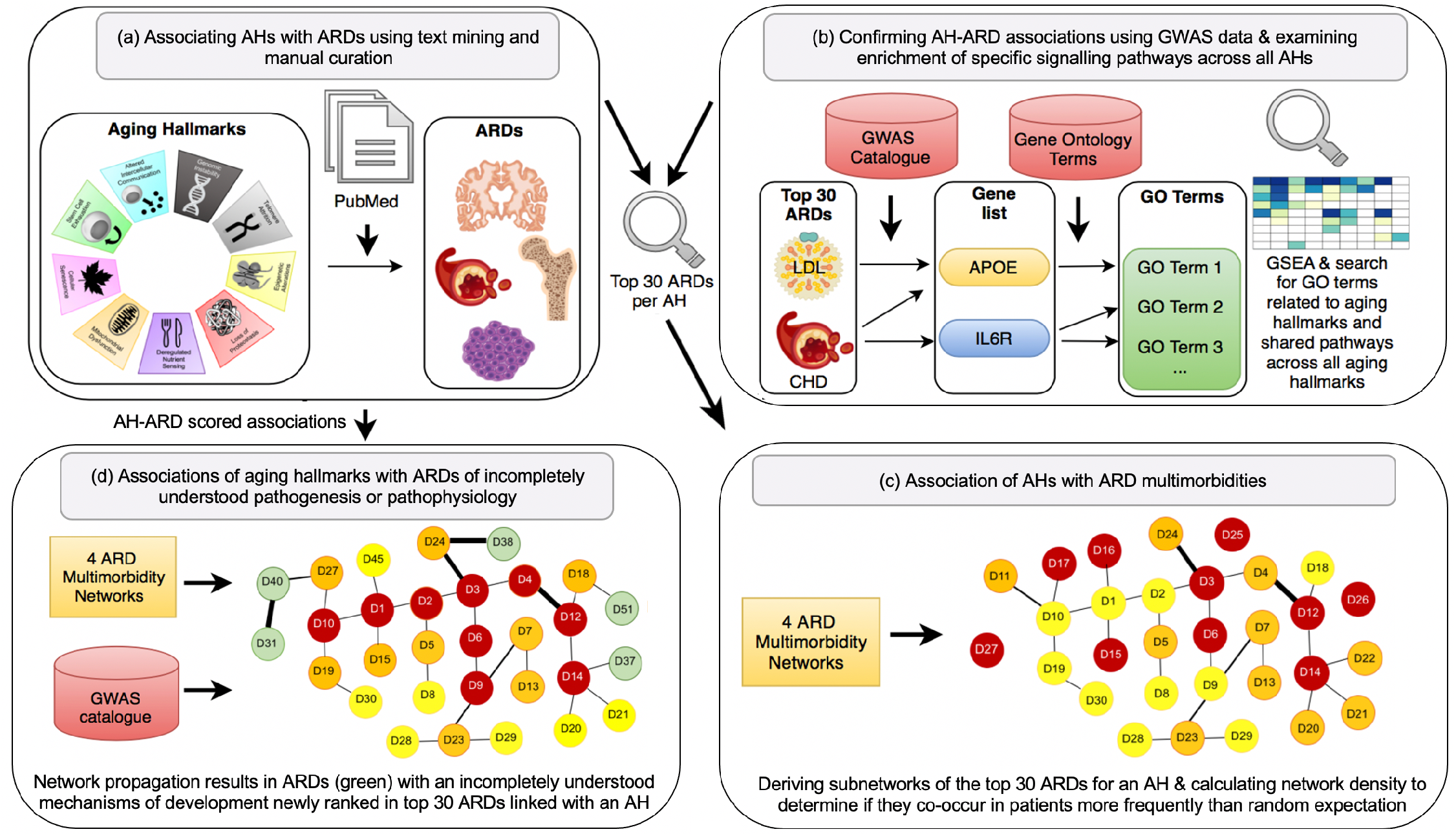
Summary of the methods. **(a) Associating aging hallmarks (AHs) with ARDs using text mining.** From 1.85 million scientific abstracts, we extracted sentences mentioning and co-mentioning aging hallmarks and ARDs to derive a score of their association. We kept scores verified using manual curation. The scores were used to identify the top 30 ranked ARDs linked to each aging hallmark. **(b) Confirming ARD-aging hallmark associations using GWAS data and investigating enrichment of specific signalling pathways across all aging hallmarks.** We identified the genes linked to each of the top 30 ARDs associated with an aging hallmark from text mining and took the union of genes, which were mapped to encoded proteins forming nine protein lists. We carried out GSEA to identify whether there was significant enrichment of GO terms related to the same aging hallmark as the ARDs were linked to in text mining. We also assessed whether there were significantly enriched signalling pathways across all aging hallmarks. **(c) Association of aging hallmarks with ARD multimorbidities**. The input data were the top 30 ARDs per aging hallmark from text mining and four ARD multimorbidity networks from age 50 years. We selected subnetworks of the top 30 ARDs per aging hallmark and compared the network density in these subnetworks to random expectation. **(d) Associations of aging hallmarks to ARDs with incompletely understood pathogenesis or pathophysiology**. We superimposed the aging hallmark-ARD scored associations from text mining onto the four ARD multimorbidity networks and iterated until convergence. We selected the top 30 ARDs based on the score of the nodes after network propagation and identified significant subnetworks. We identified ARDs with incompletely understood pathogenesis or pathophysiology newly associated with aging hallmarks (green) in the subnetworks and explored genetic data for links to the same aging hallmark.

We found strong, non-random associations between ARDs and aging hallmarks, confirmed by manual curation. This enabled us to identify the ARDs with highest evidence of association with each aging hallmark, which were verified using gene set enrichment analysis (GSEA). The genetic data also implicated roles of innate and adaptive immune, Ras-ERK, and the intrinsic apoptotic signalling pathways in the aetiology of multiple, diverse, aging-hallmark-associated ARDs. We found that ARDs with the highest evidence of association with five of the nine aging hallmarks were more likely than expected by chance to occur in patients as multimorbidities, and these associations were stable over 10-year age ranges from age 50. We also identified that aging hallmarks may provide a mechanism for the aetiology of ARDs with incompletely understood pathogenesis and/ or pathophysiology.

## Results

### Associations between aging hallmarks and ARDs in the biomedical literature

Each aging hallmark has a greater role in the development and disordered physiology of certain ARDs and a lesser role in others^1,8^. If an aging hallmark and ARD are frequently co-mentioned in the scientific literature, this association could indicate a causal connection between them. We therefore applied text mining to the biomedical literature to identify the ARDs with the highest co-mentions with each aging hallmark (Fig 2a). As the associations derived from text mining could be confounded by another factor, we verified that the aging hallmark-ARD associations derived from text mining were direct, using manual curation and confirmation from GWAS data (Fig 2b).

Our text data consisted of 1.85 million abstracts on human aging extracted from PubMed, termed the “human aging corpus”, and was separated into 20.48 million sentences (Fig 2a). Synonyms of the aging hallmarks and ARDs were needed to maximize identification of relevant sentences in the text data^20^. We therefore developed an aging hallmark taxonomy, so that synonyms and subclasses of an original aging hallmark could be brought into a dictionary for the nine aging hallmarks (Fig 1)^21^. The starting point for the aging hallmark taxonomy was *“The Hallmarks of Aging”*^1^ paper and the rationale for selection of each taxonomy term is shown in Table S1. Overall, the original nine aging hallmarks^1^ were expanded into a taxonomy of 65 related terms and four levels (Fig 1). For the development of the ARD dictionary, we used a previously developed definition, yielding a list of 207 ARDs meeting the criteria^18^ from which four ARDs that were not specific enough for scientific literature mining were excluded (Table S2). We then determined if each original aging hallmark synonym and/ or ARD synonym was mentioned in each of the 20.48 million sentences (see Methods, Fig 2a). We excluded 19 ARDs that had fewer than 250 associated sentences from abstracts in the human aging corpus (Table S2). As a co-occurrence score to quantify aging hallmark-ARD associations for the remaining 184 ARDs, we used the Ochiai coefficient^22^, which scores sentences mentioning and co-mentioning an aging hallmark and an ARD, and adjusts for uneven study density of each aging hallmark and ARD.

ARDs and aging hallmarks with higher Ochiai coefficients are likely to be related in some way, but the type of relationship, for instance a causal connection, is not known^23^. Therefore, we manually examined sentences co-mentioning aging hallmarks and ARDs for each represented aging hallmark-ARD pair to determine the type of relationship^24^. For each aging hallmark-ARD pair, we manually examined co-mentioning sentences until we had encountered a sufficient number (see Methods) that correctly reported an aging hallmark had a role in the development or disordered physiology of a disease (Table S4). Aging hallmark-ARD combinations with insufficient evidence of association from manual curation were set to zero and the Ochiai coefficient associating each aging hallmark and ARD was updated. Next, the updated Ochiai coefficients were sorted in descending order to provide a rank for association of each ARD with each aging hallmark (Fig 3a). We selected the top 30 ARDs associated with each aging hallmark based on the updated Ochiai coefficients (Fig 2a, 3b) as a sufficiently large number to explore in multimorbidity networks, while also prioritising the ARDs with the greatest literature evidence of association with an aging hallmark.

**Figure 3.**
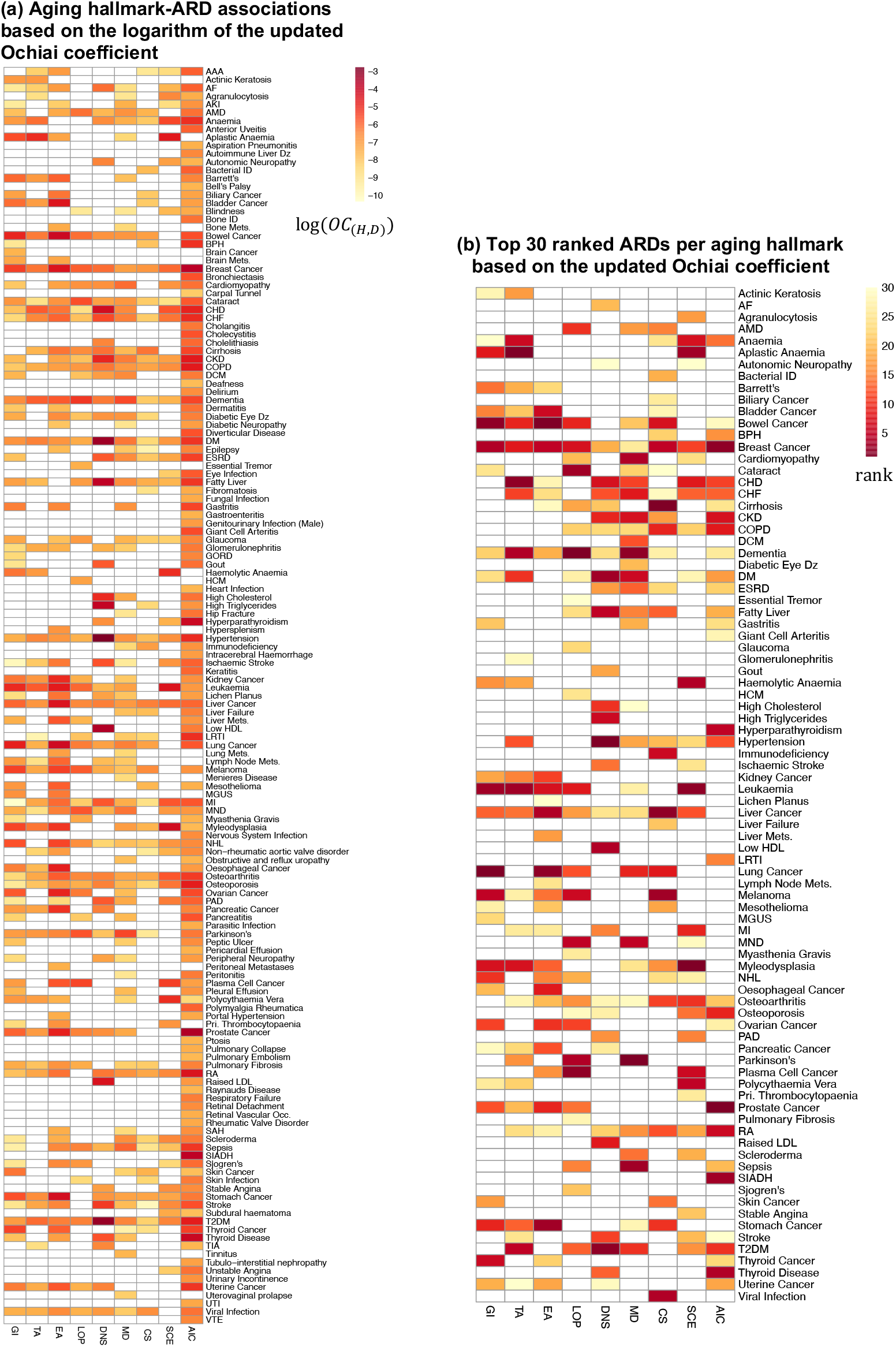
Aging hallmark-ARD associations from text mining. (a) Aging hallmark-ARD associations based on the logarithm of the updated Ochiai coefficient. The highest ranked ARDs are in red and lowest ranked in yellow. ARDs with no association are shown in white. (b) The top 30 ranked ARDs for each aging hallmark. 1st (dark red) to 30th (light yellow) ranked ARDs for a given aging hallmark are highlighted. ARDs not ranked in the top 30 are shown in white. Abbreviations: Table S9.

There were clear patterns of association between specific aging hallmarks and ARDs (Fig 3a & b). For instance, disorders frequently mentioned in association with genomic instability and epigenetic alterations were primary malignancies, such as lung cancer, bowel cancer and leukaemia (Fig 3b). This was as expected, since “genomic instability and mutation” are hallmarks of cancer and epigenetic alterations are important in cancer development and progression^10,25^. Highly ranked ARDs for telomere attrition and stem cell exhaustion were haematological disorders, including aplastic anaemia, anaemia and myelodysplasia (Fig 3b)^1^. There were strong associations between proteostasis and neurodegenerative disorders including dementia, Parkinson’s disease and motor neurone disease (MND), which are indeed associated with amyloid-beta aggregates, ∝-synuclein aggregates and dipeptide-repeat polymers, respectively (Fig 3b)^9,26^. Mitochondrial dysfunction was strongly associated with neurodegenerative disorders and cardiomyopathy, again showing that our approach could recapture established associations (Fig 3b)^8,27^. Highly ranked ARDs for cellular senescence included immunodeficiency, which is associated with immunosenescence, and cancers, which are exacerbated by the senescence-associated secretory phenotype (Fig 3b)^28,29^. Highly ranked disorders for deregulated nutrient sensing were high triglycerides, low high-density lipoprotein (HDL) cholesterol, hypertension and type 2 diabetes mellitus (T2DM) (Fig 3b). These ARDs comprise the metabolic syndrome, which is strongly associated with insulin resistance^30^. Altered intercellular communication was associated with specific malignancies and autoimmune disorders, such as prostate cancer and rheumatoid arthritis (RA), respectively (Fig 3b)^31^. Thus, our text mining approach correctly captured many molecular and cellular processes known to be involved in the respective ARD aetiology and confirmed that aging hallmark-ARD associations were highly non-random.

### Confirmation of ARD-aging hallmark associations from GWAS data

We next used genetic information to obtain confirmation of the aging hallmark-ARD associations derived from text mining. This was based on the notion that proteins encoded by genes associated with top 30 ARDs should show significant enrichment of GO terms related to the same aging hallmark on GSEA (Fig 2b). We linked the top 30 ARDs per aging hallmark to genes using the GWAS catalog^32^ (Fig 2b). We thus formed nine lists representing the union of genes linked to these top 30 ARDs (Fig 2b). As GO terms are mapped to gene products, we mapped each of the protein-coding genes to a single protein typically representing the canonical isoform, resulting in nine “protein lists” (Table 1a)^33^. We then carried out GSEA to test for significant enrichment of biological process GO terms related to the same aging hallmark (Fig 2b, Fig S1a-i). The GWAS catalog is associated with PMIDs and we avoided any risk of circularity by removing the PMIDs that intersected between studies included from the GWAS catalog and the 917,645 scientific titles/ abstracts mentioning aging hallmarks and/ or ARDs. Thus, this approach to verifying aging hallmark-ARD associations was independent of the literature-based method.

**Table 1.** Number of proteins in each aging hallmark protein list and number of proteins in each list linked to the five significant signalling pathways. We identified the genes linked to each of the top 30 ARDs associated with an aging hallmark from text mining. We took the union of genes leading to nine gene lists. Protein-coding genes within each gene list were mapped to proteins forming nine protein lists. (a) Total number of proteins in each protein list. The associated aging hallmark from text mining represents the rows in the ‘aging hallmark’ column (i.e., genomic instability (GI), telomere attrition (TA), epigenetic alterations (EA), loss of proteostasis (LOP), cellular senescence (CS), deregulated nutrient sensing (DNS), mitochondrial dysfunction (MD), stem cell exhaustion (SCE) and altered intercellular communication (AIC)). We next carried out GSEA followed by a search for GO terms mentioning “pathway” or “cascade”, which showed significant enrichment of five pathways across all aging hallmark protein lists represented in (b-f). The number of proteins in each protein list linked to the GO terms: (b) *‘IFN-γ-mediated signaling pathway’*, (c) *‘T-cell receptor signaling pathway’*, (d) *‘positive regulation of T-cell receptor signaling pathway’*, (e) *‘positive regulation of the ERK1/2 cascade’* and (f) *‘intrinsic apoptotic signaling pathways in response to DNA damage by p53 class mediator’* compared to the expected number (*p<0.05, ** p<0.01, ***p<0.001, ****p<0.0001). The ‘total’ rows show the union of proteins from all nine protein lists and the union of mapped ARDs.

| Aging hallmark | a. Total number of proteins in protein list | Number of proteins in protein list linked to signalling pathway (expected number) |  |  |  |  |
| --- | --- | --- | --- | --- | --- | --- |
| | | b. IFN-γ | c. T-cell | d. T-cell (positive regulation) | e. ERK1/2 (positive regulation) | f. intrinsic apoptotic |
| GI | 511 | 9 (2.7)*** | 13 (3.7)*** | 3 (0.4)** | 15 (6.0)** | 7 (1.4)*** |
| TA | 872 | 19 (4.7)**** | 21 (6.3)*** | 5 (0.7)*** | 27 (10.3)**** | 8 (2.4)*** |
| EA | 658 | 14 (3.5)**** | 20 (4.7)**** | 4 (0.5)** | 17 (7.8)** | 7 (1.8)*** |
| LOP | 817 | 16 (4.4)**** | 17 (5.9)*** | 4 (0.6)** | 26 (9.7)**** | 6 (2.2)* |
| DNS | 1,212 | 20 (6.5)** | 26 (8.7)**** | 4 (1.0)* | 31 (14.3)**** | 7 (3.3)* |
| MD | 1,058 | 20 (5.7)**** | 24 (7.6)*** | 5 (0.8)*** | 31 (12.5)**** | 8 (2.9)** |
| CS | 594 | 10 (3.2)** | 17 (4.3)*** | 3 (0.5)** | 16 (7.0)** | 9 (1.6)**** |
| SCE | 680 | 17 (3.7)**** | 17 (4.9)** | 4 (0.5)** | 23 (8.0)**** | 7 (1.8)*** |
| AIC | 809 | 14 (4.3)*** | 19 (5.8)*** | 3 (0.6)* | 24 (9.6)**** | 7 (2.2)** |
| Total (union of encoded proteins) |  | 25 | 30 | 5 | 40 | 9 |
| Total (union of mapped ARDs) |  | 21 | 19 | 9 | 22 | 11 |

We next tested whether biological processes related to each aging hallmark were indeed significantly enriched in the protein list representing the top 30 ARDs associated with that hallmark (Fig 2b, Fig S1a-i). Between 511 and 1,212 proteins were associated with each of the aging hallmarks (Table 1a). We carried out GSEA and searched for GO terms related to each aging hallmark (Fig S1a-i). We identified significant enrichment of terms related to the same aging hallmark as was associated to the ARDs via text mining (Fig S1a-i). For example, “DNA damage response”, “telomere maintenance”, “regulation of autophagy”, “replicative senescence”, “glucose homeostasis”, “regulation of mitochondrion fission” and “stem cell differentiation” were significantly enriched in the genomic instability, telomere attrition, loss of proteostasis, cellular senescence, deregulated nutrient sensing, mitochondrial dysfunction and stem cell exhaustion protein lists, respectively (Fig S1a, b, d-h). The altered intercellular communication protein list showed significant enrichment of processes related to hormone synthesis and inflammatory response while the epigenetic alterations protein list showed significant enrichment of terms related to histone acetylation (Fig S1c, i). Thus, the protein lists derived from the aging hallmark-associated gene lists were significantly enriched for annotations related to their own aging hallmark. Therefore, analysis of GWAS data confirmed the specific associations between aging hallmarks and ARDs that had been found from the literature co-occurrence scores (Fig 2a, b).

### Enrichment of signalling pathways across all aging hallmarks

Our literature mining revealed highly specific associations between ARDs and aging hallmarks, and these were independently confirmed by GWAS data. However, hallmarks of aging are part of a complex nexus of failure of molecular and cellular processes, are not independent of each other, and may share some common underlying signalling pathways. Therefore, we explored whether common signalling pathways were shared across all aging hallmark protein lists and, thus, contribute to the development of multiple aging hallmark-associated ARDs. Strikingly, for the ARDs that were associated with specific hallmarks and that were present in our GWAS analysis, there was clear evidence from the GWAS data for commonalities in the signalling cascades and pathways across all aging hallmark protein lists (Fig 4a). GSEA followed by search for GO terms mentioning “pathway” or “cascade” showed that five pathways were significantly enriched in all aging hallmark protein lists (Fig 4a, Table 1b-f). Three were linked to the innate and adaptive immune system, including the “interferon-γ-mediated signaling pathway” and the “T-cell receptor signaling pathway” and to its “positive regulation” (Fig 4a, Table 1b-d). These pathways are interconnected, as interferon-γ is a cytokine produced by multiple immune cells including cells of the adaptive immune system, such as T-cells^34^. “Positive regulation of the ERK1/2 cascade” and the “intrinsic apoptotic signalling pathway in response to DNA damage by a p53 class mediator” were also significantly enriched across all aging hallmark protein lists (Fig 4a, Table 1e, f).

**Figure 4.**
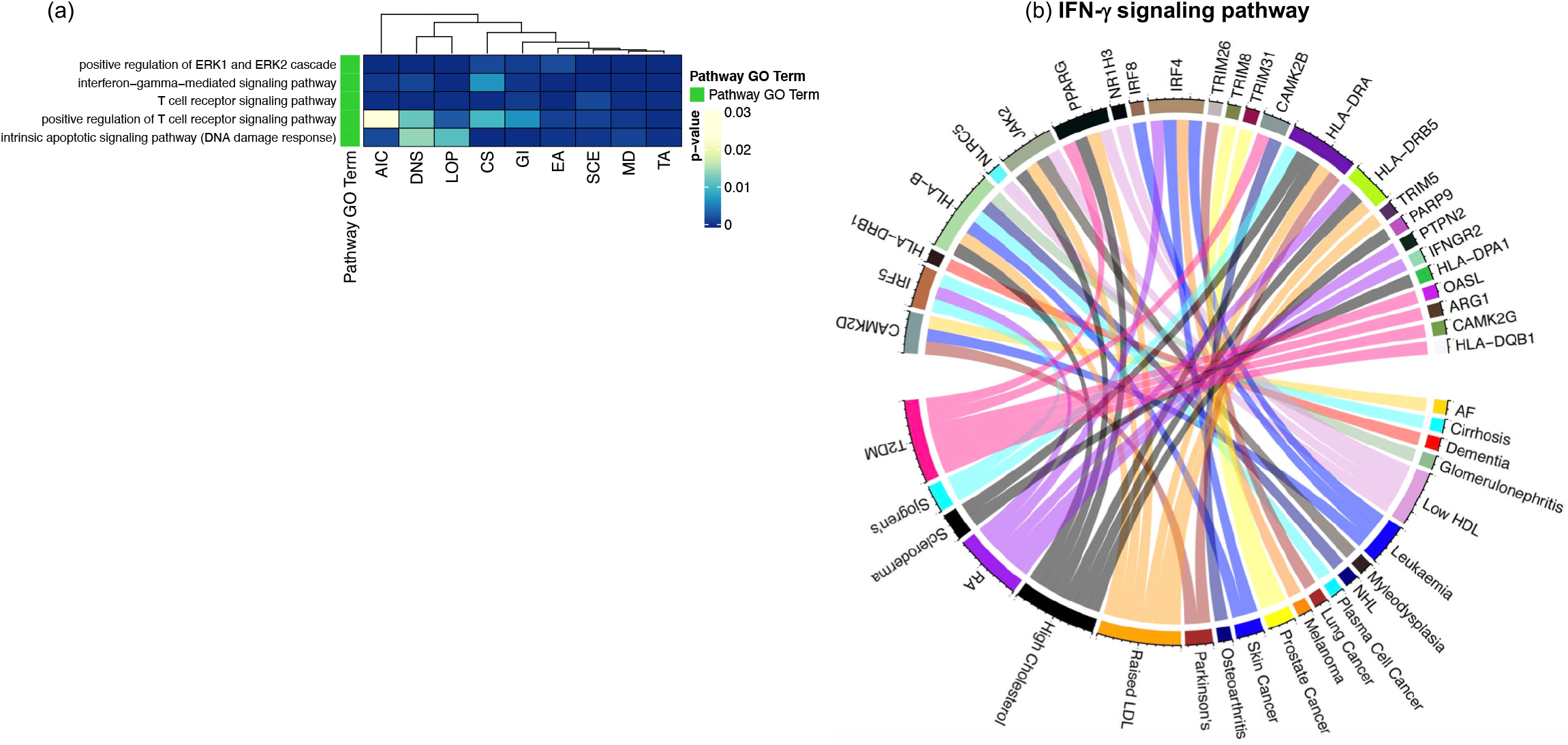
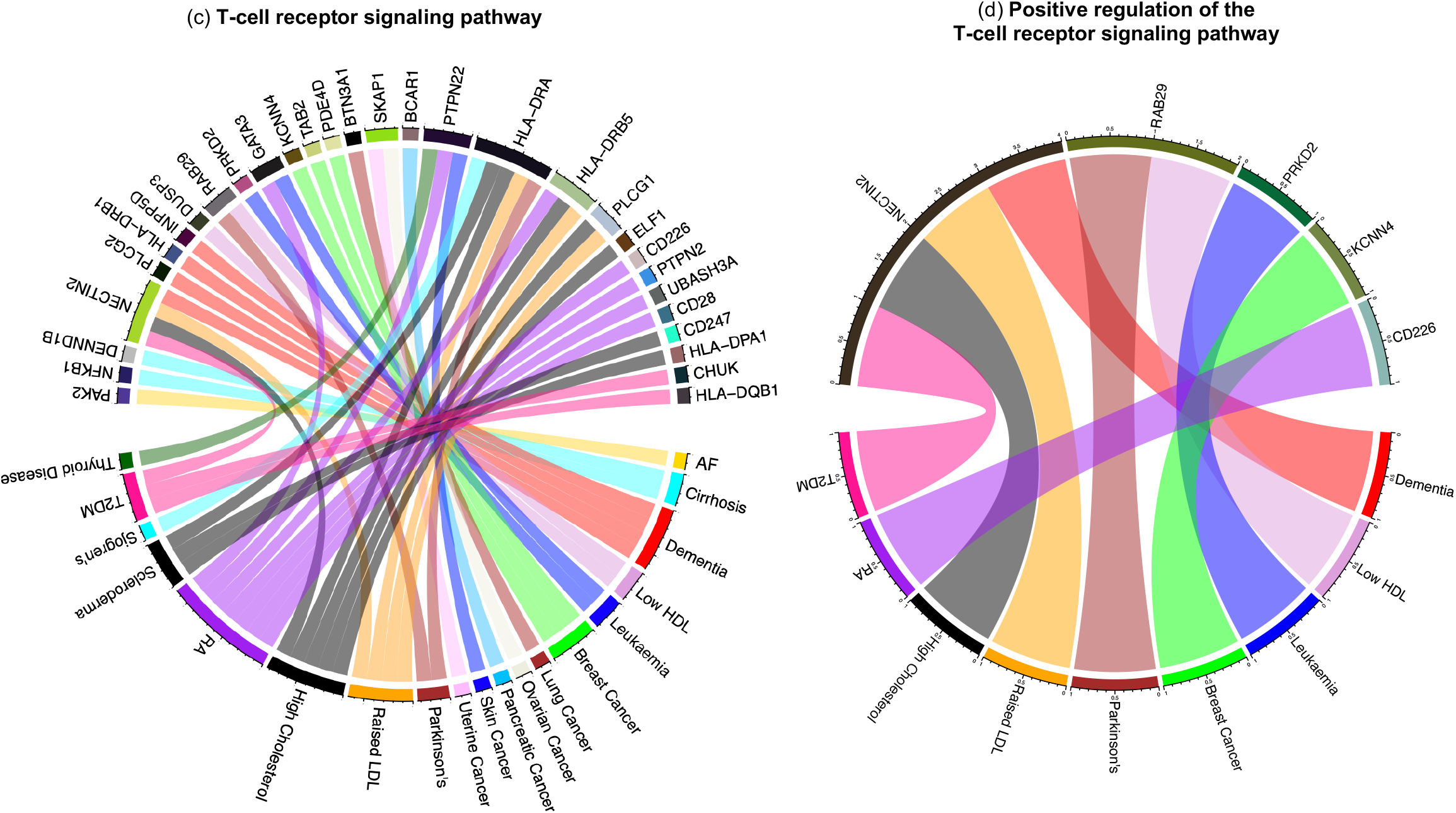
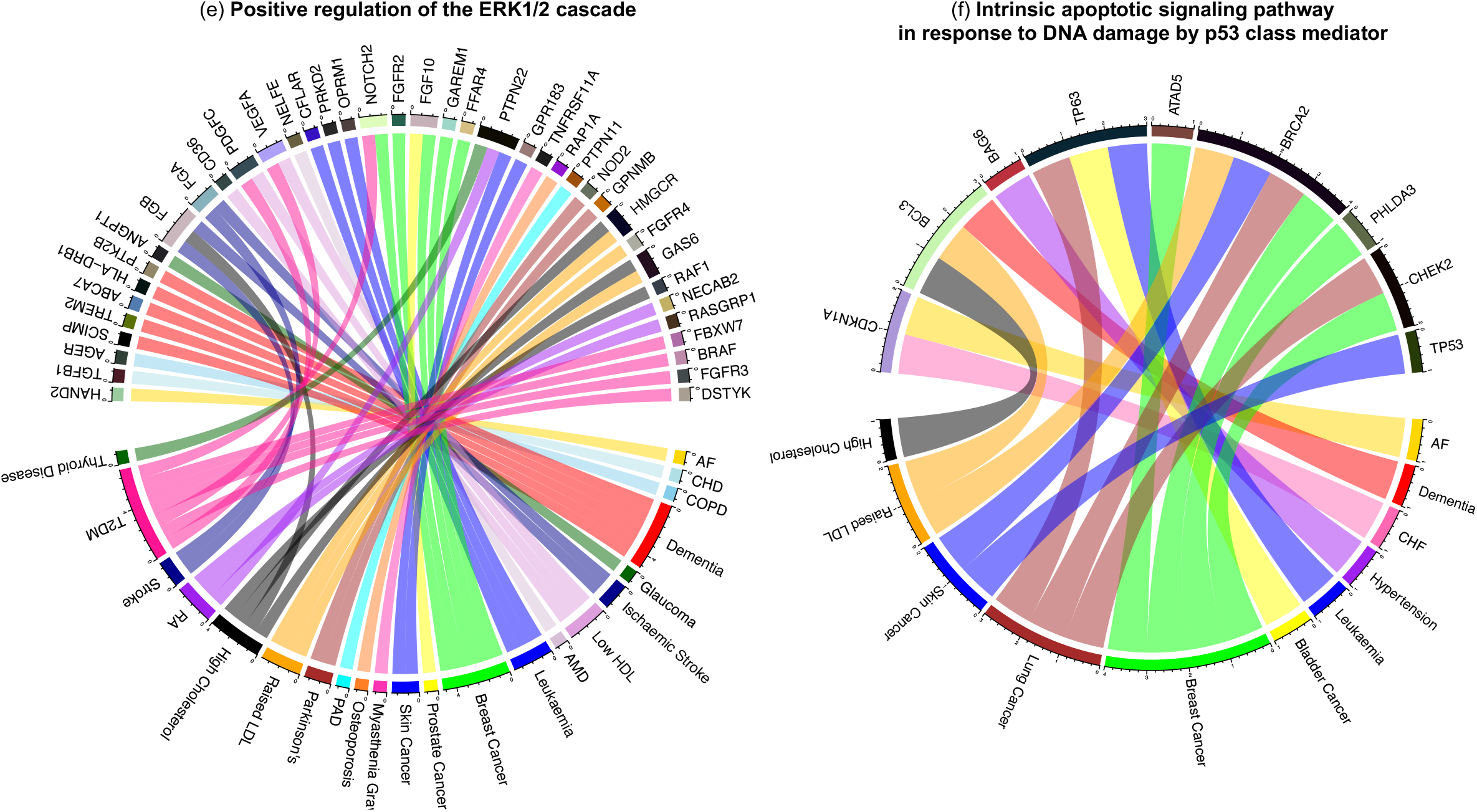
Significantly enriched signalling pathways across all aging hallmark protein lists. (a) P-values of enriched signalling pathways across all aging hallmarks. We identified the genes linked to each of the top 30 ARDs associated with an aging hallmark from text mining and took the union of genes. These were mapped to encoded proteins forming nine protein lists. The associated aging hallmark from text mining represents the column labels of the heatmap. We carried out GSEA and searched for GO terms related to signalling pathways. Five signalling pathways were significantly enriched across all aging hallmark protein lists. (b-f) The union of proteins/ genes associated with each of the five significantly enriched pathways was derived and they were linked to their associated ARDs. These are shown in the circos plots representing: (b) IFN-γ-mediated signalling pathway, (c) T-cell receptor signalling pathway, (d) positive regulation of T-cell receptor signalling pathway, (e) positive regulation of the ERK1/2 cascade, and (f) the intrinsic apoptotic signalling pathway in response to DNA damage by p53 class mediator. Abbreviations: Table S9.

To explore these common pathways further, we derived the union of proteins associated with each of the GO terms across all aging hallmarks, mapped them to their underlying genes and linked them to their associated ARDs (Table 1b-f). A total of 21 ARDs were linked to 25 genes encoding proteins associated with the interferon-γ pathway (Fig 4b, Table 1b), 19 to 30 genes encoding proteins associated with the T-cell receptor signalling pathway (Fig 4c, Table 1c), 9 to 5 genes encoding proteins associated with positive regulation of the T-cell receptor signalling pathway (Fig 4d, Table 1d), 22 to 40 genes encoding proteins associated with the ERK1/2 cascade (Fig 4e, Table 1e) and 11 to 9 genes encoding proteins associated with the intrinsic apoptotic signalling pathway (Fig 4f, Table 1f). These signalling cascades are therefore implicated in the aetiology of these diverse, aging-hallmark-associated ARDs.

### Association of aging hallmarks with ARD multimorbidities

The results of our literature mining revealed specific associations between aging hallmarks and ARDs, and these were independently confirmed by GWAS data. We next explored the possible role of aging hallmarks in the co-occurrence of two ARDs in the same patient, known as multimorbidity (Fig 2c). To do this, we assessed whether ARDs associated with the same aging hallmark occurred more frequently in the same patient than random pairs of ARDs. We used previously created multimorbidity networks^19^ reflecting non-random co-occurrence of two diseases in the same patient. The multimorbidity networks were created for different age classes by binning electronic health records of 3.01 million individuals into nine 10-year age intervals^18,19,35^. Within each age interval, significantly co-occurring disease pairs were linked in the respective network (see Methods)^19^. The stratification by age accounts for the fact that occurrence^35^ and co-occurrence^19^ of diseases change with age. Since we were particularly interested in ARDs, we used the four networks for the age groups of 50 years and over for subsequent analyses because 170 of the 184 ARDs had a median age of onset ≥ 50 years (Fig 2c)^18^. Thereby, we obtained four networks of 184 ARDs (Table S6)^18,19^.

We assessed whether the ARDs associated with each aging hallmark were more likely to occur as multimorbidities than expected by chance. If multimorbidities were at least partially explained by common underlying hallmarks, we would expect that ARDs targeted by the same hallmark should be more tightly connected in the multimorbidity networks. To test that notion, we selected the top 30 ARDs for each aging hallmark and extracted the subnetworks consisting of those 30 diseases (Fig 2c; Fig 3b), resulting in 36 subnetworks for the four age-specific ARD multimorbidity networks and the nine aging hallmarks. A higher observed network density than expected by chance indicates that there are more edges than expected, and hence that the ARDs within the subnetwork are more frequently multimorbidities than random ARD sets of the same size. To calculate the p-value, we compared the observed network density of the subnetwork with the network density expected by chance using permutation tests.

For five of nine aging hallmarks, namely deregulated nutrient sensing (p < 0.0001), mitochondrial dysfunction (p < 0.05), cellular senescence (p < 0.05), stem cell exhaustion (p < 0.001) and altered intercellular communication (p < 0.01), the nodes representing the top 30 associated ARDs were connected by more edges than expected by chance across all age categories (Table 2, Fig 2c). The ARDs associated with these five aging hallmarks thus co-occurred in individual patients more frequently than expected by chance and these associations were stable over 10-year age ranges from age 50 years (Fig 5a-e, Table 2). For example, the deregulated nutrient sensing multimorbidity subnetwork contained nodes connected by edges representing the progression of known multimorbidities, such as type 2 diabetes mellitus with fatty liver (Fig 5a)^36^. These non-random associations suggest that these five aging hallmarks do indeed have a role in the development of ARD multimorbidity in patients (Table 2).

**Figure 5.**
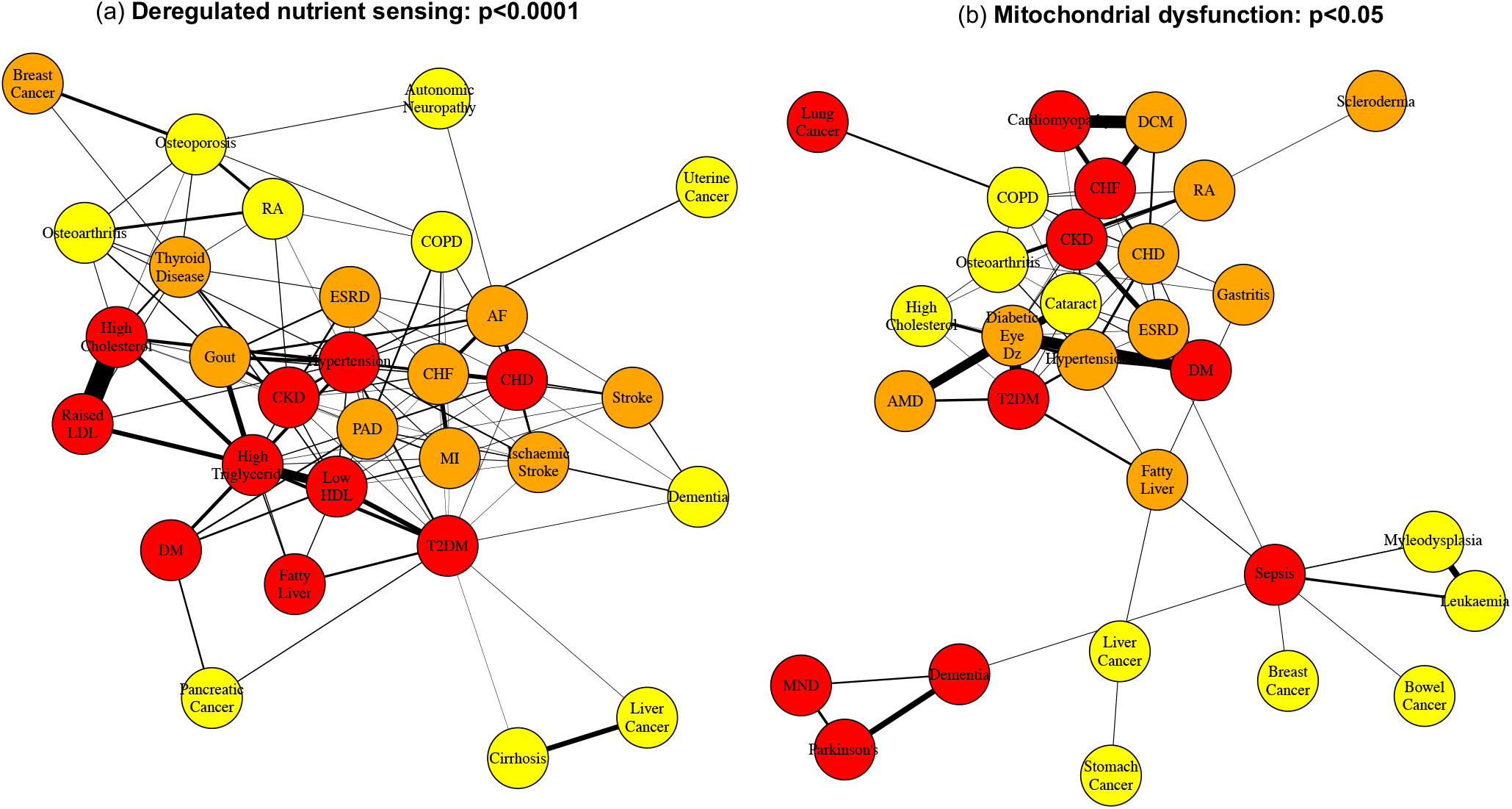
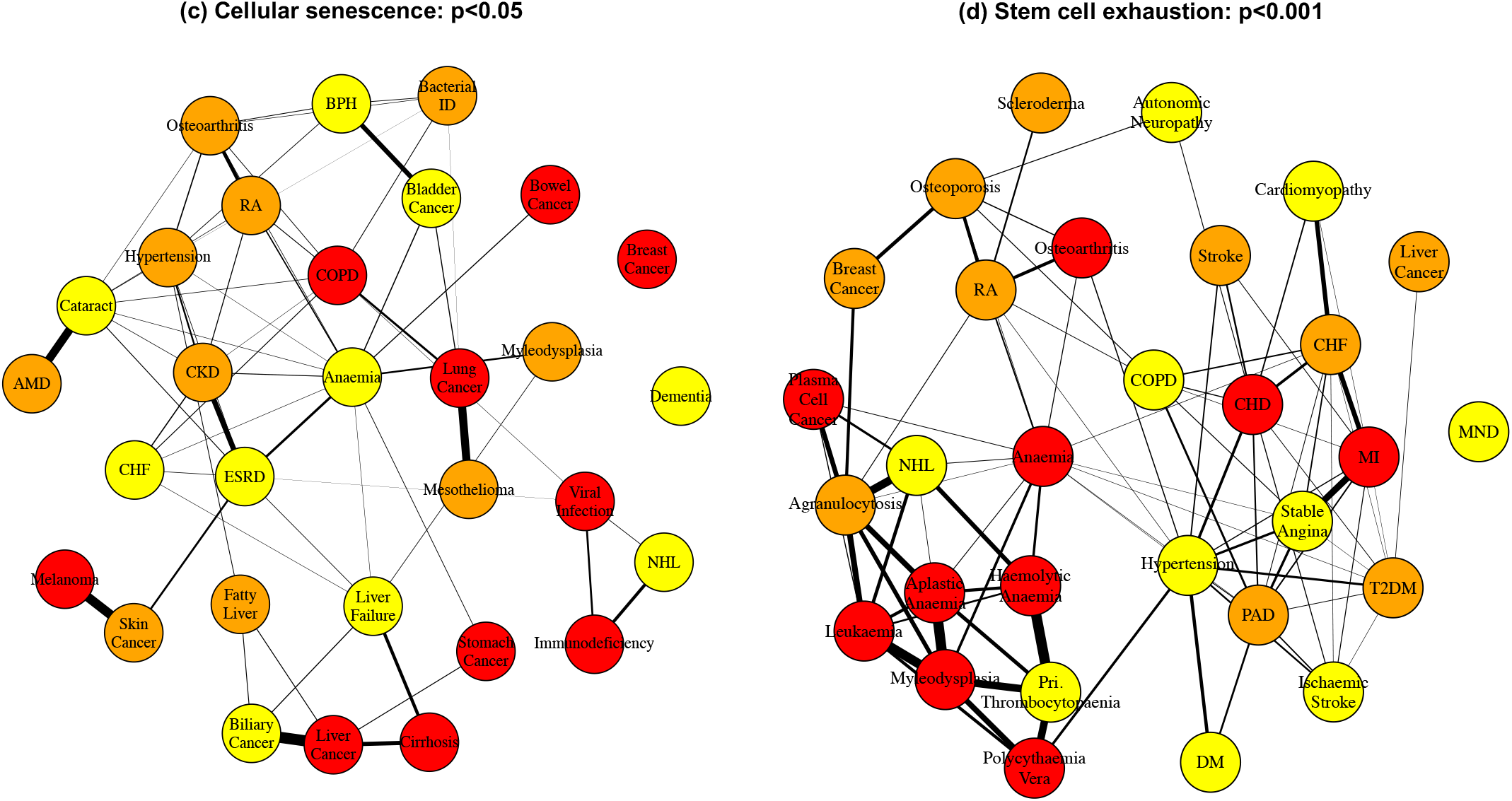
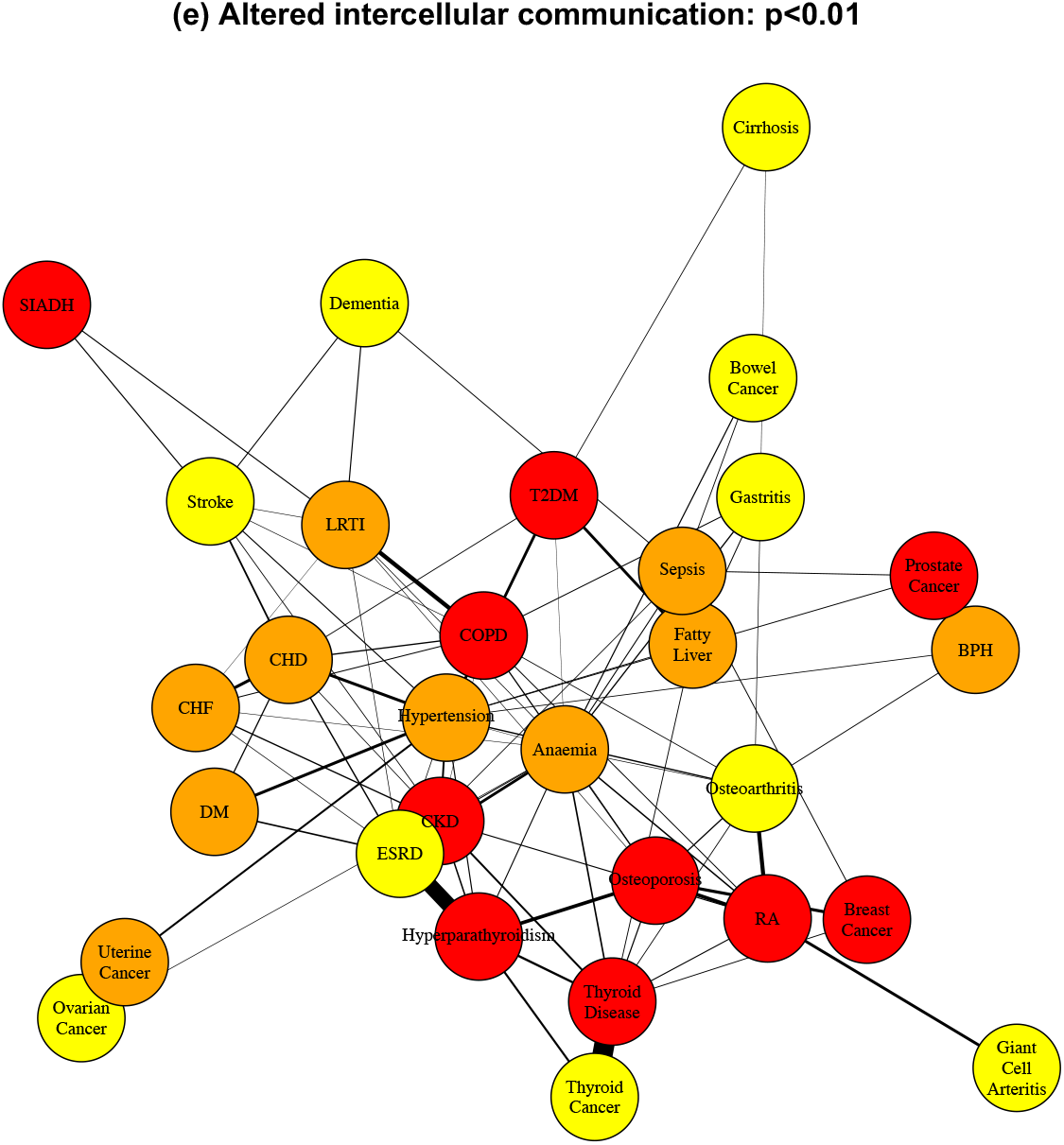
Subnetworks containing nodes representing the top 30 ranked ARDs for each aging hallmark (50-59 year age category). The (a) deregulated nutrient sensing, (b) mitochondrial dysfunction, (c) cellular senescence, (d) stem cell exhaustion, and (e) altered intercellular communication subnetworks. Nodes are coloured by ARD ranking for a given aging hallmark: the 1^st^ to 10^th^ ranked in red, the 11^th^ to 20^th^ ranked in orange and the 21^st^ to 30^th^ ranked in yellow. Abbreviations: Table S9.

**Table 2.**
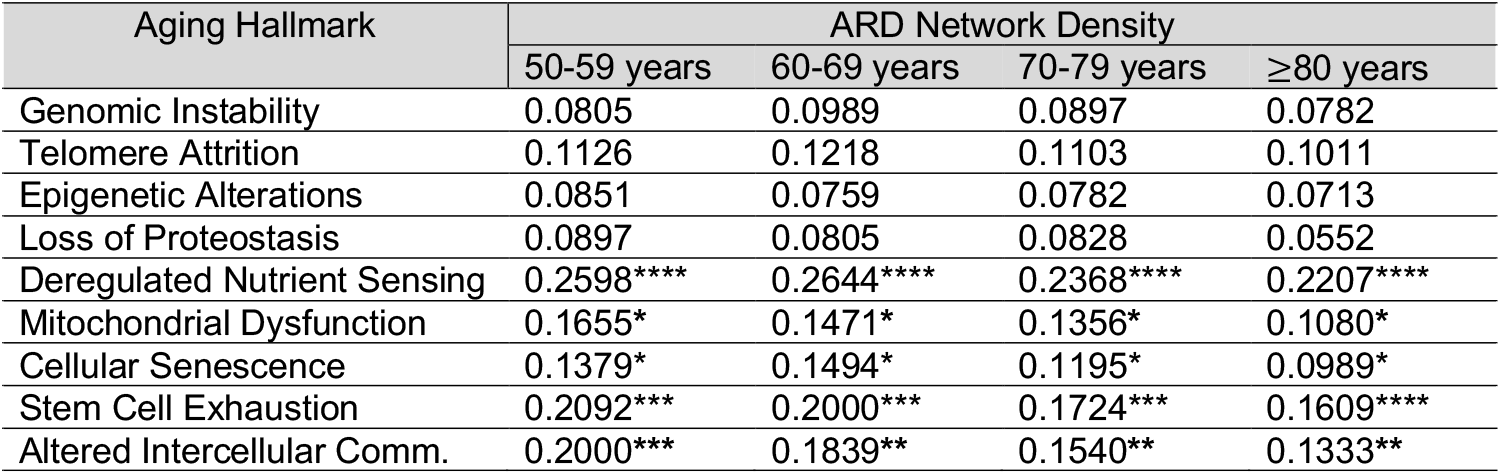
Network density of subnetworks of the top 30 ranking ARD nodes compared to random expectation for age categories 50-59 years, 60-69 years, 70-79 years and ≥80 years. The number of times the network density from permutations (n = 20,000) was greater than or equal to the true network density for that aging hallmark was used to calculate the p-value. The p-value was corrected for multiple testing across the 4 age categories per aging hallmark using the Benjamini-Hochberg procedure (*p < 0.05, ** p <0.01, *** p <0.001, ****p<0.0001).

| Aging Hallmark | ARD Network Density |  |  |  |
| --- | --- | --- | --- | --- |
| | 50-59 years | 60-69 years | 70-79 years | $\geq 80$ years |
| Genomic Instability | 0.0805 | 0.0989 | 0.0897 | 0.0782 |
| Telomere Attrition | 0.1126 | 0.1218 | 0.1103 | 0.1011 |
| Epigenetic Alterations | 0.0851 | 0.0759 | 0.0782 | 0.0713 |
| Loss of Proteostasis | 0.0897 | 0.0805 | 0.0828 | 0.0552 |
| Deregulated Nutrient Sensing | 0.2598**** | 0.2644**** | 0.2368**** | 0.2207**** |
| Mitochondrial Dysfunction | 0.1655* | 0.1471* | 0.1356* | 0.1080* |
| Cellular Senescence | 0.1379* | 0.1494* | 0.1195* | 0.0989* |
| Stem Cell Exhaustion | 0.2092*** | 0.2000*** | 0.1724*** | 0.1609**** |
| Altered Intercellular Comm. | 0.2000*** | 0.1839** | 0.1540** | 0.1333** |

### Associations of aging hallmarks with ARDs with incompletely understood pathogenesis or pathophysiology

The analysis above suggests that ARDs that are tightly connected in the multimorbidity networks are more likely affected by the same hallmark than random pairs of diseases. Thus, we speculated that this association could be used to identify hallmark-ARD associations that were so far unknown, i.e. based on the fact that many neighbouring ARDs in the network are associated with a common hallmark (‘guilt by association’)^37^. Therefore, we focused on ARDs with incompletely understood pathogenesis or pathophysiology, that were not originally ranked in the top 30 ARDs associated with a hallmark, but where the hallmark may nonetheless contribute to aetiology.

For each aging hallmark, we superimposed the aging hallmark-ARD cooccurrence scores (or updated Ochiai coefficients) from text mining onto the respective ARD nodes in each of the four multimorbidity networks (Fig 2d). The scores were then smoothed over the network, which amplifies regions where ARDs have higher co-occurrence scores with a given aging hallmark and dampens regions with lower scores^37^ and thus assigns relatively high scores to ARDs that are surrounded in the network by ARDs associated with a common hallmark. Since this process changes the ARD-hallmark associations of all diseases in the network, it also changes the ranking of ARDs associated with each aging hallmark (Fig 2d). We identified those subnetworks with a significantly greater network density than random expectation and identified newly prioritized ARDs within them (Table S7).

Two ARDs with incompletely understood mechanism of pathogenesis or pathophysiology were newly ranked among the top 30 ARDs, namely essential tremor and Bell’s Palsy (Table S7, Fig S2a & b)^38,39^. Essential tremor is a neurological disorder characterised by an involuntary, rhythmic tremor and was newly prioritized as a top 30 ARD associated with mitochondrial dysfunction (Fig S2a). It has previously been associated with mitochondrial abnormalities; however, the degree of their role is unclear^40^. This disorder also has genetic evidence of association with five genes (i.e., STK32B, NAT2, LINGO1, CTNNA3 and LRRTM3) at genome wide significance. However, we cannot exclude that the association is a consequence of initial misdiagnosis, such as of Parkinson’s disease as essential tremor^41^. Bell’s palsy was newly prioritized as a top 30 ARD associated with deregulated nutrient sensing, which has previously been reported to be associated with prognosis of the Bell’s palsy^42^. However, the association may also be a consequence of initial misdiagnosis of diabetic mononeuropathy as Bell’s palsy (Fig S2b)^43^. There were no reported genetic associations with Bell’s palsy in the GWAS catalog at genome wide significance. Overall, our findings indicate that aging hallmarks may contribute to a better understanding of disease aetiology.

## Discussion

The contribution of aging hallmarks to development of ARD multimorbidities in humans is largely unexplored. We have combined aging hallmark-ARD associations derived from text mining, verified using genetic data, with disease networks derived from electronic health records, to explore whether mechanisms of aging contribute to ARD multimorbidities in humans, and might therefore be targeted to prevent or treat more than one ARD simultaneously.

First, using two independent approaches, we explored patterns of association between specific aging hallmarks and ARDs. We text mined 917,645 literature abstracts, followed by manual curation, and found strong, non-random associations between ARDs and aging hallmarks. We verified the associations that emerged by GSEA using GO annotations of proteins encoded by genes linked to the top 30 ARDs. By integrating our findings with networks of multimorbidities, we found that five aging hallmarks were indeed associated with human ARD multimorbidities. Deregulated nutrient sensing, mitochondrial dysfunction, cellular senescence, stem cell exhaustion and altered intercellular communication, were associated with the co-occurrence of ARDs in individual patients more than expected by chance. Reassuringly, these aging hallmarks were associated with ARD multimorbidity across all four decadal age-ranges, and the associations were often highly significant. Overall, these findings indicate that therapies targeted at each of these five aging hallmarks may prove to be beneficial in the prevention of the associated ARD multimorbidities in humans. For instance, sirolimus and related compounds inhibit the TORC1 complex in the nutrient-sensing network^44^, and can both extend healthy lifespan in model organisms^45^ and boost the response to vaccination against influenza in elderly people^46^. Senolytics and senescence associated secretory phenotype (SASP) modulators eliminate senescent cells and inhibit the SASP respectively, and thus target the cellular senescence hallmark^28^, and can both improve tissue health during aging and increase lifespan in mice^47^ and may prevent cellular senescence-associated ARD multimorbidities^48^.

In model organisms, targeting common signalling pathways delays the onset of ARDs and extends lifespan^1,15–17^. Specific signalling pathways are intertwined with the aging hallmarks, for example, the IIS pathway is associated with the deregulated nutrient sensing aging hallmark^1^. Aging hallmarks are not independent of each other with, for instance, DNA damage and telomere shortening contributing to cellular senescence^49^ and loss of stem cell function^50^. Thus, aging hallmarks may share some common underlying pathways, which also contribute to the development of multiple aging hallmark-associated ARDs. Strikingly, five signalling pathways/ cascades were significantly enriched across the protein lists for all nine aging hallmarks. These pathways are therefore likely to play a key role in the aetiology of ARDs. Among these five signalling pathways, three were involved in the innate and/ or adaptive immune response. The underlying genes were derived from ARDs comprising metabolic syndrome disorders, auto-immune disorders and cancers, thus highlighting the importance of the immune response across multiple ARDs^11^. The “intrinsic apoptotic signalling pathway in response to DNA damage by a p53 class mediator” was also significantly enriched across all aging hallmark protein lists. The underlying genes were derived from multiple cancers and metabolic syndrome disorders^10,51^.The ERK1/2 pathway regulates many processes including cell survival, metabolism and inflammation^52^ and was significantly enriched across all aging hallmark protein lists. The underlying genes were derived from 22 aging hallmark-associated ARDs (Fig 4e) and indeed activation of the ERK1/2 pathway has been suggested to play a role in these ARDs either directly or through their risk factors. For example, increased activity of the ERK1/2 pathway has been identified in type 2 diabetes mellitus^53^ and hypertension^54^, which are major risk factors for cardiovascular disorders. Additionally, activating mutations upstream of ERK1/2 contribute to over fifty percent of human cancers^55^. Increased phosphorylation of cellular ERKs has also been identified in the thyroid disorder, hypothyroidism^56^, and in atrial fibrillation^57^. Furthermore, ERK1/2 inhibition reduces beta-amyloid neurotoxicity in Alzheimer’s disease^52^, decreases inflammation and apoptosis in stroke patients^52^ and prevents rheumatoid arthritis in mouse models^58^. Interestingly, the ERK1/2 cascade is linked to aging in model organisms and the MEK inhibitor, Trametinib, prolongs lifespan in *Drosophila*^15^. Thus our analysis suggests that inhibition of the ERK1/2 pathway could prevent up to 22 human aging hallmark-associated ARDs.

Using network propagation, we identified ARDs with incompletely understood pathogenesis where aging hallmarks may contribute to their development. Essential tremor has previously been associated with mitochondrial abnormalities, but the degree of their role is unclear^39,40^. We found that essential tremor co-occurred with many ARDs strongly linked to mitochondrial dysfunction implying this is in fact a key pathogenic mechanism in essential tremor. However, we cannot exclude the association is a consequence of initial misdiagnosis, such as of Parkinson’s disease as essential tremor^41^. Our findings were also supported by genetic data, as essential tremor is also associated with the variant N-acetyltransferase 2 (NAT2) gene. NAT2 is associated with insulin resistance^59^, and deficiency of the mouse orthologue (i.e., NAT1) has also been associated with mitochondrial dysfunction^60^. Therefore, aging hallmarks may contribute to the development of ARDs with incompletely understood mechanisms of development.

A potential limitation is that, because certain ARDs occupy more of the scientific research effort, there is a risk that they would be more frequently included in the top 30 ARDs associated with aging hallmark and, therefore, included in multimorbidity subnetworks. We used four approaches to reduce the risk of this, which were (*i*) adjusting for uneven study density on each ARD by using a cooccurrence score, (*ii*) excluding ARDs with less than 250 associated sentences from abstracts in the human aging corpus, (*iii*) keeping only aging hallmark-ARD associations verified by manual curation and (*iv*) verifying aging hallmark-ARD associations using genetic data. An additional potential limitation is that ARD multimorbidities may be connected in electronic health records due to incorrect initial diagnosis, which may complicate the evaluation of incompletely explained ARDs. These limitations will be overcome as our knowledge of the aging hallmarks, ARD multimorbidities and genes underlying ARDs improves.

Our study provides evidence for the role of aging hallmarks in the aetiology of human ARD multimorbidities and ARDs with incompletely understood pathogenesis. We also raise the possibility that multiple ARDs may be prevented by targeting common signalling pathways, such as the innate and adaptive immune pathways, the intrinsic apoptotic signalling pathway and the ERK1/2 pathway. Future work will determine whether a prophylactic agent or cure for human ARD multimorbidities can be developed by targeting each of five aging hallmarks.

## Methods

The methods are summarised in Figure 2.

### Information retrieval of the ‘human aging corpus’

A set of primary research articles (or corpus) on human aging was required for text mining. Our corpus was developed by defining inclusion and exclusion criteria followed by retrieving 1.93 million PubMed identifiers (PMIDs) of abstracts meeting those criteria from PubMed (Table S8a, b). The 1.93 PMIDs representing title/ abstracts on human aging meeting our search criteria were retrieved from the PubMed database using the Biopython Entrez application programming interface^61^ on 10^th^ April 2020. Next, the 2019 PubMed baseline contains over 29 million abstracts and was downloaded in Extensible Mark-up Language (XML) format. Data was extracted from the XML files to produce separate, comma-separated values (CSV) files containing 29,138,919 million rows and six columns including titles, abstracts and PMIDs. The rows containing the 1.93 million PMIDs of the human aging corpus were identified. PMIDs associated with missing data were eliminated and, subsequently, the text data was cleaned. This gave 1.85 million abstracts representing the “human aging corpus”, which were tokenized into 20.48 million sentences.

### Information extraction by dictionary-based methods with co-occurrence scoring

#### Aging hallmark dictionary

An aging hallmark taxonomy was developed to maximize retrieval of relevant literature on each aging hallmark from PubMed (Fig 1). We modelled our methodology on the approach used previously to develop a cancer hallmarks taxonomy^10,21^. The starting point for the taxonomy was the original *“The Hallmarks of Aging”*^1^ paper from which we selected relevant subcategories of the nine original aging hallmarks; however, occasionally, we inferred a particular taxonomy term that was not specifically stated in original paper (Fig 1, Table S1). Additional taxonomy levels represented increasingly specific biological processes within a subclass (Table S1). Synonyms for each aging hallmark taxonomy term were retrieved from the Unified Medical Language System (UMLS) Metathesaurus^62^ from the U.S. National Library of Medicine (NLM) and relevant review articles. The aging hallmark taxonomy term synonyms were combined to form an aging hallmark dictionary and then linked to the corresponding original aging hallmarks.

#### Age-related disease dictionary

The ARD definition was developed previously by applying a hierarchical agglomerative clustering algorithm to clinical data on 278 diseases^18^. Four of nine “main” clusters contained 207 diseases and these diseases also had an adjusted R^2^ of greater than 0.85 on the Gompertz-Makeham model^18^. These 207 diseases were defined as ARDs (Table S2)^18^. Four ARDs that did not translate effectively to scientific text mining approaches were eliminated from further analysis (Table S2). We next retrieved synonyms for each of the remaining 203 ARDs from the MeSH thesaurus from the NLM^63^. The Comparative Toxicogenomics Database’s ‘merged disease vocabulary’^64^ was downloaded on 21^st^ March 2019. It contains the MeSH diseases hierarchy processed in a CSV file. Supplementary concepts and animal diseases were excluded. This left 4,789 human diseases mapped to 28,638 entry terms, or synonyms, after processing. MeSH terms were assigned to the 188 of 203 ARDs from the 4,789 diseases. The 188 ARDs were mapped to a hierarchical tree of 1,427 rows containing MeSH term subclasses of assigned MeSH terms, of which, 545 relevant subclasses were kept. The synonyms to each subclass were edited manually and then merged for each ARD. For the remaining 15 ARDs, synonyms were derived from the Unified Medical Language System (UMLS) Metathesaurus^62^. The synonyms were merged to form an ARD dictionary and then linked to the corresponding 203 ARDs.

#### Calculating the Ochiai coefficient

The aging hallmark dictionary and human ARD dictionary were matched against the 20.48 million sentences from PubMed titles and abstracts. 19 ARDs with less than 250 associated sentences were eliminated (Table S2). The co-occurrence of the nine aging hallmarks with the remaining 184 ARDs was scored at the sentence level using the Ochiai coefficient^22^ (Equation 1). The Ochiai coefficient (*OC*_(*H,D*)_) adjusts for the fact that certain ARDs are frequently studied in the biomedical literature while others are infrequently studied. For a given aging hallmark and ARD, nHD is the total number of sentences that co-mention the aging hallmark and ARD. nD and nH are the total number of sentences that mention the ARD and aging hallmark, respectively (Equation 1)^65^.

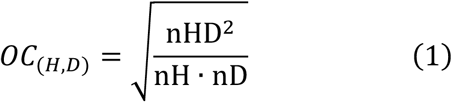

#### Verifying extracted associations by manual curation

ARDs and aging hallmarks with higher Ochiai coefficients are likely to be related in some way, but the type of relationship is not known^23^. Therefore, we manually assessed the sentences co-mentioning aging hallmarks and ARDs to determine whether they correctly reported an association between the aging hallmark and ARD (Table S5)^24^. Our approach to manual curation was to define co-mentioning sentences as either (1) “confirmed association” where an aging hallmark is reported (or inferred) to have a role in the ARD development or persistence, (2) “no association”, (3) “irrelevant” or (4) “error”^66^(Table S3). For aging hallmarks with less than 2,500 co-mentioning sentences, we manually examined all sentences co-mentioning a given aging hallmark-ARD pair until we found one sentence that satisfied the criteria of “confirmed association” (Table S3 & S5). For the remaining aging hallmarks, three sentences that satisfied the criteria of “confirmed association” were required (Table S3 & S5). If an aging hallmark-ARD pair could not be confirmed by a sufficient number of sentences, its Ochiai coefficient was set to zero to increase the reliability of our findings. The 30 highest scoring ARDs were selected for each aging hallmark.

### Analysis of aging hallmark-associated multimorbidity subnetworks and network propagation

#### Multimorbidity Networks

We used multimorbidity networks derived from previously analysed clinical data on 289 diseases, including the 184 ARDs, in 3.01 million individuals^18,19^. The clinical data was obtained from Clinical Practice Research Datalink (CPRD), which was linked to the Hospital Episode Statistics admitted patient care (HES APC) dataset and accessed via the CALIBER research platform^18,19^. From the multimorbidity network data, we derived an undirected ARD network, where the nodes represent the 184 ARDs which were connected by edges. Edges linked ARD nodes if they were linked by a positive, significant partial correlation (after Bonferroni correction). The partial correlation was used as the edge weight^18,19^. 170 of the 184 ARDs had a median age of first recorded diagnosis 50 years or older^18,19^. Therefore, we used four multimorbidity networks for the 184 ARDs representing age categories from 50 years (Table S6)^18,19^.

#### Network analysis of top 30 ranked aging hallmark-associated ARDs

We selected the top 30 ranking nodes for each aging hallmark from each of the four multimorbidity networks and, therefore, plotted 36 subgraphs. The network density (*D*) was calculated for each subnetwork using the algorithm shown in Equation 2 where *E* is the number of edges in a subnetwork and *V* is the number of nodes in a subnetwork.

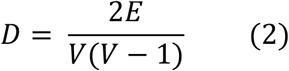

For each aging hallmark and age category, we shuffled the updated Ochiai coefficient associated with the 184 ARDs 20,000 times. At each shuffle, we selected the top 30 ARD nodes to form a subnetwork and calculated their network density. For a given permutation, each time the random network density (*D_k_*) was greater than or equal to the actual network density (*D_o_*) we added a score of 1, and otherwise 0. The p-value (*p*) for the network density was derived using Equation 3 where *K* is the total number of permutations^67^.

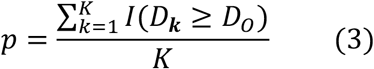

The p-value was corrected for multiple testing across the 4 age categories per aging hallmark using the Benjamini-Hochberg procedure^68^.

#### Network propagation onto multimorbidity networks

For each aging hallmark and age category, the updated Ochiai coefficient scores (*F*^0^) were superimposed onto each of the ARD nodes of the multimorbidity network. Using a Random Walk with Restart (RWR) algorithm the scores were smoothed over the network (Equation 4) from the R package BioNetSmooth version 1.0.0 to derive the posterior score^69^.

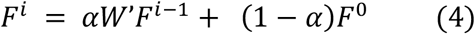

In the RWR algorithm, F^*i*^ and F^*i*−1^ are the posterior evidence of association of an aging hallmark with an ARD at smoothing iteration, *i* and *i* −1, respectively, and we iterated until convergence (*i* = 30). The degree row-normalized adjacency matrix of the weighted disease network is represented by *W′*. The entries in the adjacency matrix (i.e. *W′* = [*w′′*_*r,c*_]) are defined in Equation 5,

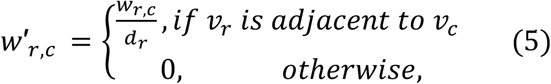

where *d_r_* is the degree of the ARD node *v_r_* and the edge weight between ARD node *v_r_* and ARD node *v_c_* is *w_r,c_*. Alpha (*α*) was set at 0.5. The top 30 ARDs with the highest posterior score after network propagation were selected to form a subnetwork. Significant subnetworks were identified using the approach described previously (Equation 3) with correction for multiple testing^68^. We identified ARDs newly prioritized in the top 30 ARDs associated with an aging hallmark in these subnetworks, which also had an incompletely understood pathogenesis or pathophysiology.

### Identification of functionally enriched biological processes using genetic data

#### Genes underlying ARDs

The NHGRI-EBI GWAS Catalog^32^ was downloaded on 26^th^ February 2020. 103 of the 203 defined ARDs were represented in the GWAS catalog^32^. These 103 ARDs were mapped to 181 ‘Mapped Traits’, which are terms from the Experimental Factor Ontology that are assigned to each GWAS and represent, for example, the disease investigated^32^. Single nucleotide polymorphisms (SNPs) with a p-value of < 5 x 10^-8^ associating them to ARDs were kept. GWAS studies in European populations were included; however, certain groups were excluded (e.g., Amish). SNPs located were assigned to genes (i.e., Ensembl Gene IDs) if they were located within a gene or intergenic SNPs less than 50 kilobase pairs (kbp) from a gene. For newly prioritized ARDs after network propagation, intergenic SNPs were assigned to genes at a distance of 75 kbp to maximize retrieval of relevant genes. The Ensembl gene IDs were mapped to National Centre for Biotechnology Information (NCBI) Gene IDs, where available, using the NCBI Gene database of *Homo sapiens*^70^. Thus, 2,364 NCBI Gene IDs were linked to 84 ARDs and 135 Mapped Traits subclasses.

#### Functional enrichment of biological processes for top 30 ARDs mapped to aging hallmarks

We identified the union of genes linked to the top 30 ARDs per aging hallmark (based on text mining) (Fig 2b). The NCBI Gene IDs for protein-coding genes were mapped to *“stringId”*s using the STRING database forming nine protein lists^33^. Of all 86 ARDs included in top 30 ranked node subnetworks, 55 were associated with 1,698 NCBI Gene IDs and mapped to 1,693 stringId. The background set was also downloaded from the STRING database on 27^th^ January 2019^33^, which contained 16,598 stringIds mapped to the biological process GO terms. 1,560 of 1,693 stringIds were also in the background set^11^. We used topGO^71^ with the Fisher’s exact test to identify biologically enriched processes against the background set and applied the ‘weight01’ algorithm to reduce redundancy of GO terms. The final p-value cut-off was 0.05 and the minimum node size was 5. Using our previously created aging hallmark dictionary, we searched for GO terms related to the aging hallmarks. Shortened synonyms and abbreviations were appended to the dictionary for specific aging hallmarks. We also searched for GO terms related to “pathway” and “cascade” and we kept only the pathways that were significantly enriched across all aging hallmark protein lists.

### Computational analyses & images

Computational analyses were carried out in Python 3.7.0 and R Version 3.3.0 and 3.6.0. Aging hallmark and ARD images were downloaded from Adobe Stock and Shutterstock after obtaining a standard license.

## Acknowledgements

HCF (Helen C Fraser) is supported by a PhD studentship from the UK Medical Research Council (grant number MR/N013867/1). LP (Linda Partridge) is supported by a Wellcome Trust Strategic Award and the Max Planck Society. AB (Andreas Beyer) is supported by the German Federal Ministry of Education and Research (BMBF; grant number HiGHmed 01ZZ1802U). VK (Valerie Kuan) is supported by the Wellcome Trust (WT 110284/Z/15/Z) and the Dunhill Medical Trust (RPGF1806/67). ADH (Aroon D Hingorani) is an NIHR Senior Investigator and supported by BHF Accelerator Award AA/18/6/34223. RJ (Ronja Johnen) is supported by the German Research Foundation (CRC1310, project number 325931972). MZ (Magdalena Zwierzyna) was supported by BenevolentAI. We thank JG (Jan Grossbach) for providing advice on gene set enrichment analysis and AL (Andre Lopes) for providing advice on statistical analysis.

## Author Contributions

Conceptualization and design of the study: HCF, LP, AB, RJ and VK. Performed the analysis: HCF. Interpreted the data and the results: HCF, LP, AB, RJ, VK, MZ, ADH, AL and JG. Drafted the paper and reviewed the drafts: HCF, LP, AB, VK, MZ, RJ, ADH.

## Data and code availability statement

The custom code to reproduce the analysis, and the data sets generated and analysed in this research article are available on request.

## Supporting information

Table S1-S9 and Figures S1-S2 can be found in the attached file.

## Ethical compliance

The multimorbidity networks were created previously using clinical data^18,19^. The clinical data was obtained from the CPRD, which was linked to HES APC dataset. These studies were approved Independent Scientific Advisory Committee (ISAC) for the Medicines and Healthcare products Regulatory Agency (MHRA) under “protocol 16_022”. At the time of data collection, participant consent was obtained and does not need to be repeated for every study.

