## Supplementary Figures and Tables for "Biological mechanisms of aging predict age-related disease multimorbidities in patients"

**Figure S1**. **P-value of enriched biological processes across all aging hallmarks.** We identified the genes linked to each of the top 30 ARDs associated with an aging hallmark from text mining. We took the union of genes leading to nine gene sets. Protein-coding genes within each gene set were mapped to proteins forming nine protein sets. The associated aging hallmark from text mining represents the column labels (i.e. GI, TA, EA, LOP, CS, DNS, MD, SCE and AIC). We carried out GSEA and expected significant enrichment of GO terms related to the same aging hallmark to verify our findings from text mining. After GSEA, we searched for GO terms related to aging hallmarks: (a) GI, (b) TA, (c) EA, (d) LOP, (e) DNS, (f) MD, (g) CS, (h) SCE and (i) AIC. The 'aging hallmark (AH) GO Term' column highlights the retrieved GO terms linked to each aging hallmark. The abbreviations are listed in Table S9.

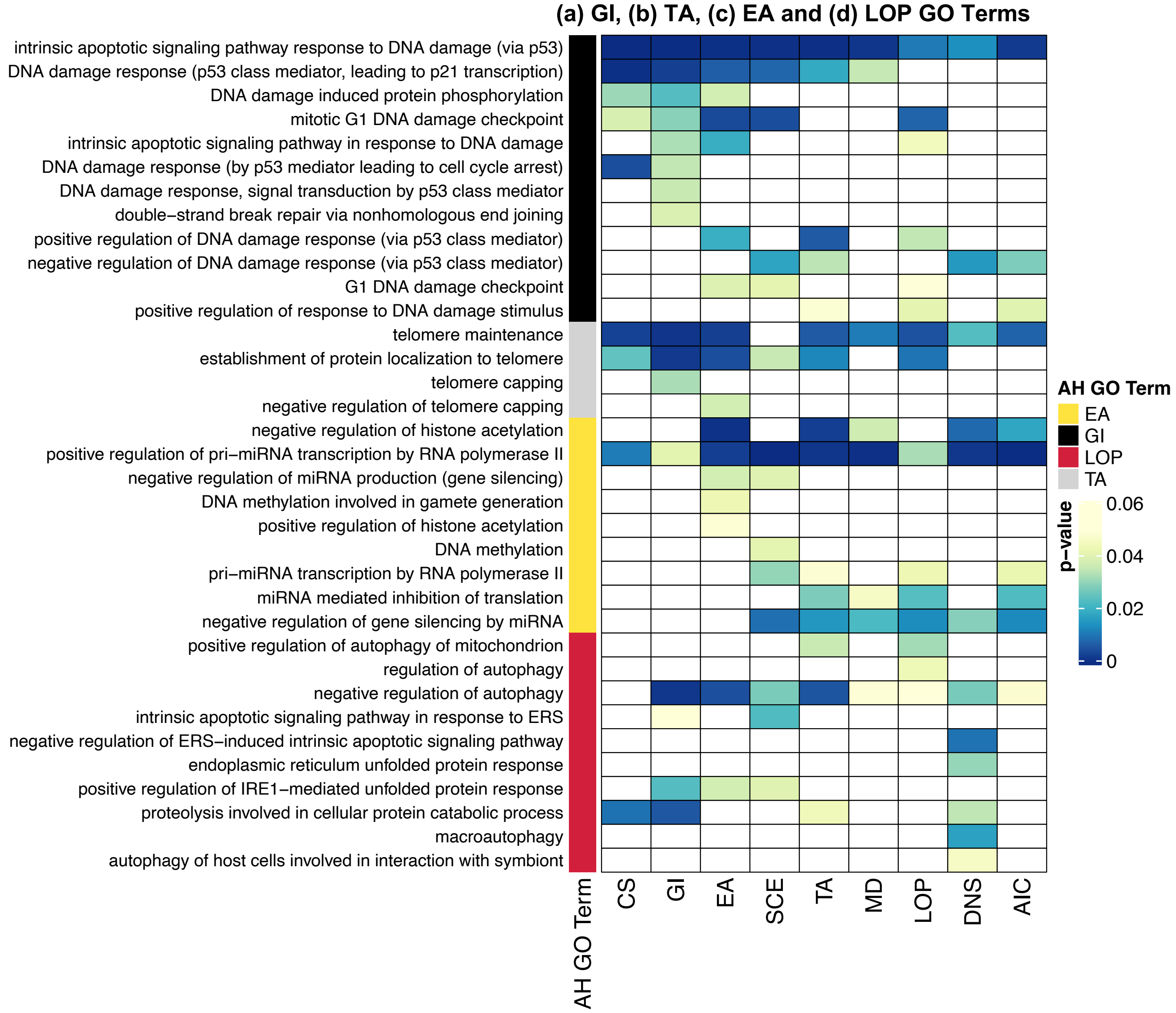

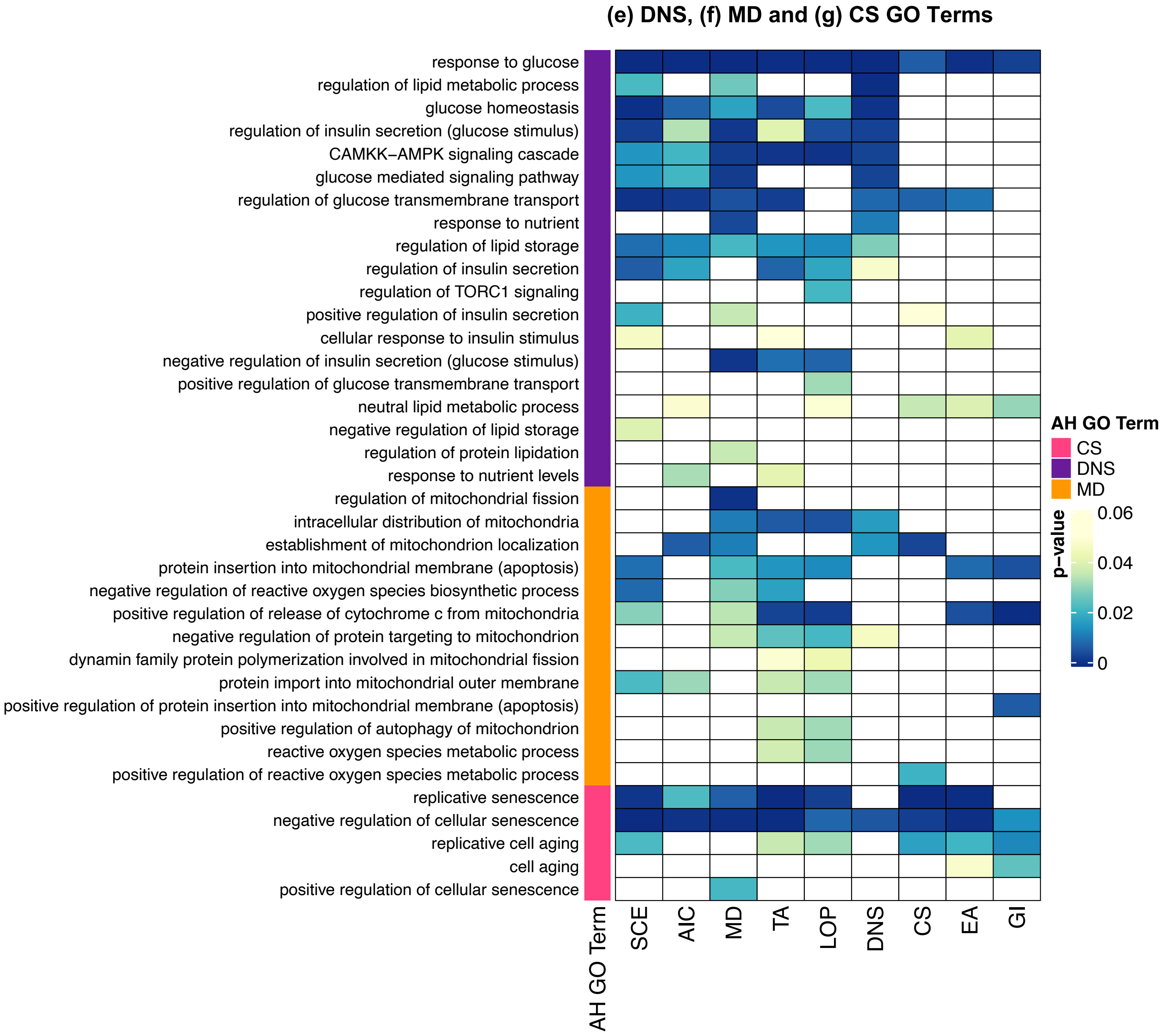
(e) DNS, (f) MD and (g) CS GO Terms
response to glucose
regulation of lipid metabolic process
glucose homeostasis
regulation of insulin secretion (glucose stimulus)
CAMKK−AMPK signaling cascade
glucose mediated signaling pathway
regulation of glucose transmembrane transport
response to nutrient
regulation of lipid storage
regulation of insulin secretion
regulation of TORC1 signaling
positive regulation of insulin secretion
cellular response to insulin stimulus
negative regulation of insulin secretion (glucose stimulus)
positive regulation of glucose transmembrane transport
neutral lipid metabolic process
negative regulation of lipid storage
regulation of protein lipidation
response to nutrient levels
regulation of mitochondrial fission
intracellular distribution of mitochondria
establishment of mitochondrion localization
protein insertion into mitochondrial membrane (apoptosis)
negative regulation of reactive oxygen species biosynthetic process
positive regulation of release of cytochrome c from mitochondria
negative regulation of protein targeting to mitochondrion
dynamin family protein polymerization involved in mitochondrial fission
protein import into mitochondrial outer membrane
positive regulation of protein insertion into mitochondrial membrane (apoptosis)
positive regulation of autophagy of mitochondrion
reactive oxygen species metabolic process
positive regulation of reactive oxygen species metabolic process
replicative senescence
negative regulation of cellular senescence
replicative cell aging
cell aging
positive regulation of cellular senescence
AH GO Term
SCE
AIC
MD
TA
LOP
DNS
CS
EA
GI
AH GO Term
CS
DNS
MD
p−value
0.06
0.04
0.02
0

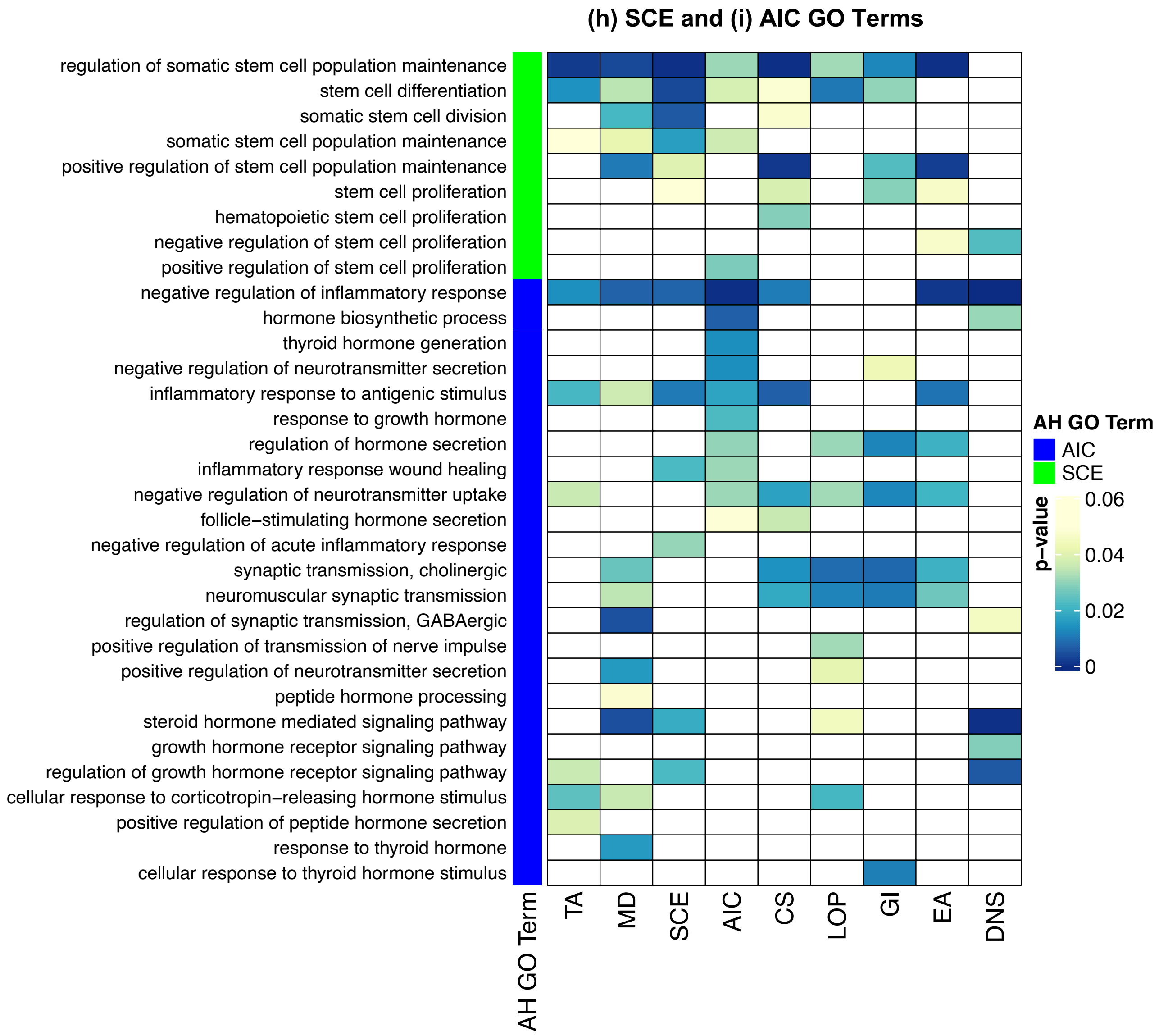

(h) SCE and (i) AIC GO Terms
regulation of somatic stem cell population maintenance
stem cell differentiation
somatic stem cell division
somatic stem cell population maintenance
positive regulation of stem cell population maintenance
stem cell proliferation
hematopoietic stem cell proliferation
negative regulation of stem cell proliferation
positive regulation of stem cell proliferation
negative regulation of inflammatory response
hormone biosynthetic process
thyroid hormone generation
negative regulation of neurotransmitter secretion
inflammatory response to antigenic stimulus
response to growth hormone
regulation of hormone secretion
inflammatory response wound healing
negative regulation of neurotransmitter uptake
follicle−stimulating hormone secretion
negative regulation of acute inflammatory response
synaptic transmission, cholinergic
neuromuscular synaptic transmission
regulation of synaptic transmission, GABAergic
positive regulation of transmission of nerve impulse
positive regulation of neurotransmitter secretion
peptide hormone processing
steroid hormone mediated signaling pathway
growth hormone receptor signaling pathway
regulation of growth hormone receptor signaling pathway
cellular response to corticotropin−releasing hormone stimulus
positive regulation of peptide hormone secretion
response to thyroid hormone
cellular response to thyroid hormone stimulus
AH GO Term
TA
MD
SCE
AIC
CS
LOP
GI
EA
DNS
AH GO Term
AIC
SCE
p−value
0.06
0.04
0.02
0

**Figure S2. Subnetworks of top 30 ranked ARDs after network propagation for (a) mitochondrial dysfunction (60-69 years) and (b) deregulated nutrient sensing (60-69 years).** Nodes are coloured by ARD ranking after network propagation for a given aging hallmark. The 1st to 10th ranked ARDs for a given aging hallmark are red, the 11th to 20th ranked ARD for a given aging hallmark are in orange and the 21st to 30th ranked ARDs for a given aging hallmark are yellow. The abbreviations are listed in Table S9.

(a) **Mitochondrial dysfunction (60-69 years): p <0.001**

(b) **Deregulated nutrient sensing (60-69 years): p<0.0001**

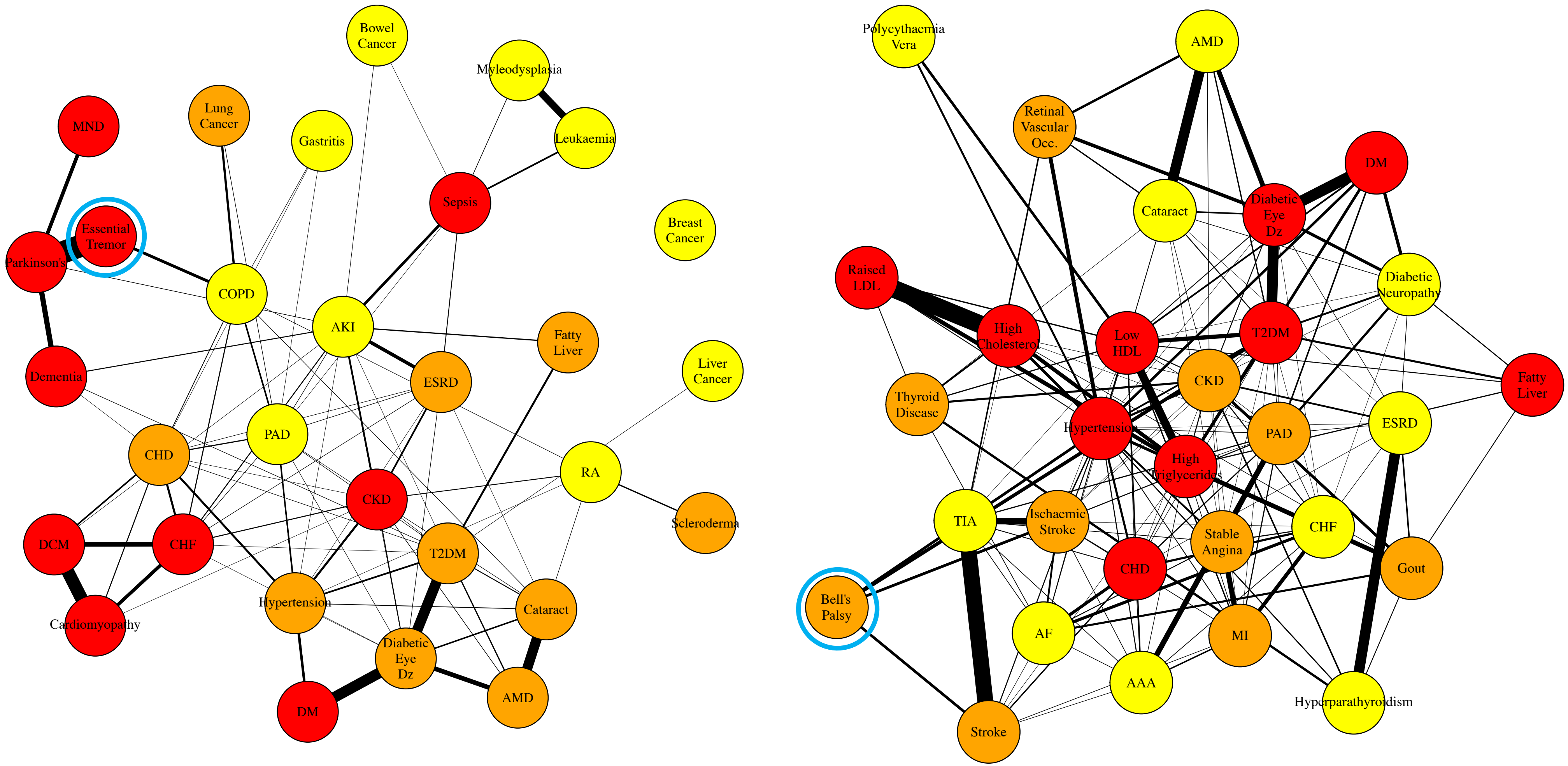

### Supplementary Tables

**Table S1. The 65 aging hallmark taxonomy terms derived from "*The Hallmarks of Aging*" paper.** The table shows quotations from "*The Hallmarks of Aging*" paper by Lopez-Otin *et al*. (2013), which support the selection of taxonomy terms. Occasionally, taxonomy terms are inferred rather than being directly mentioned in "*The Hallmarks of Aging*" , which is indicated with (*i*).

| Original Aging Hallmark | Taxonomy Term(s) | Supporting quotations from "*The Hallmarks of Aging*" (Lopez-Otin *et al*., 2013) or rationale |
|---|---|---|
| **Genomic Instability** | Genomic instability | "**Genomic Instability**" (Hallmark of Aging) |
| | Somatic mutations | "**Somatic mutations** accumulate within cells from aged humans." |
| | DNA damage,<br>Chromosomal breakage | "Other forms of **DNA damage**, such as **chromosomal aneuploidies** and copy number variations, have also been found associated with aging." |
| | Transposable elements | "The genetic lesions arising from extrinsic or intrinsic damage are highly diverse and include … gene disruption caused by the integration of viruses or **transposons**". |
| | DNA repair deficiencies | "**Deficiencies in DNA repair mechanisms** cause accelerated aging." |
| | mtDNA mutations | "**Mutations** and deletions in aged **mtDNA** may also contribute to aging." |
| | mtDNA damage | "The first evidence that **mtDNA damage** might be important for aging and ARDs derived from the identification of human multisystem disorders caused by mtDNA mutations." |
| | Double strand (ds)-DNA breaks | "Note that both nuclear DNA and mitochondrial DNA are subjected to age-associated genomic alterations … **double strand breaks**." |
| | DNA breaks (*i*),<br>Single strand (ss)-DNA breaks (*i*) | "The **genetic lesions** arising from intrinsic and extrinsic damages are highly diverse and include point mutations … ". Additional examples include **single-strand DNA breaks** or, more generally, **DNA breaks**[1]. |
| **Telomere Attrition** | Telomere attrition | "**Telomere Attrition**" (Hallmark of Aging) |
| | Decreased telomere length,<br>Decreased leukocyte telomere length (*i*) | "In humans, recent meta-analyses have indicated a strong relation between **short telomeres** and mortality risk, particularly at younger ages." **Leukocyte telomere length** is a more specific subcategory of decreased telomere length[2]. |
| **Epigenetic Alterations** | Epigenetic alterations | "**Epigenetic Alterations**" (Hallmark of Aging) |
| | Gene transcription,<br>coding-RNAs | "Aging is associated with an increase in **transcriptional** noise, and an **aberrant production and maturation of many mRNAs**." |
| | microRNAs,<br>non-coding RNAs | "The aging-associated transcriptional signatures also affect **non-coding RNAs**, including a class of **miRNAs** that is associated with the aging process". |
| | Histone modifications,<br>histone acetylation,<br>histone methylation,<br>DNA methylation | "Alterations in the **acetylation** and **methylation** of **DNA** or **histones**, as well as of other chromatin-associated proteins, can induce epigenetic changes that contribute to the aging process." |

| | | |
|---|---|---|
| **Loss of Proteostasis** | Loss of proteostasis | "**Loss of Proteostasis**" (Hallmark of Aging) |
| | Proteolysis, autophagy, proteasome | "The activities of the two principal **proteolytic** systems implicated in protein quality control, namely, the **autophagy-lysosomal** system and the **ubiquitin-proteasome system**, decline with aging." |
| | Protein aggregation | "Failure to refold or degrade unfolded proteins can lead to their accumulation and **aggregation**, resulting in proteotoxic effects." |
| | ER stress, unfolded protein response (UPR) | "Endogenous and exogenous stress causes the unfolding of proteins … **ER Stress. Unfolded proteins** are usually refolded by heat-shock proteins (HSP) or targeted to destruction by the ubiquitin-proteasome or lysosomal (autophagic) pathways." |
| | Chaperone | "The stress-induced synthesis of cytosolic and organelle-specific **chaperones** is significantly impaired in aging." |
| **Deregulated Nutrient Sensing** | Deregulated nutrient sensing | "**Deregulated Nutrient Sensing**" (Hallmark of Aging) |
| | Insulin resistance, dyslipidaemia *(i)* | "Thus, a **decreased IIS** is a common characteristic of both physiological and accelerated aging, while a constitutively decreased IIS extends longevity." **Dyslipidaemia** is also associated with deregulated nutrient sensing[3]. |
| | IIS pathway, nutrient-sensing pathways | "In addition to the **IIS pathway** that participates in glucose-sensing, three additional related and interconnected **nutrient-sensing systems** are the focus of intense investigation…" |
| | mTORC1 | "These observations, together with those involving the IIS pathway, indicate that intense trophic and anabolic activity, signalled through the IIS or the **mTORC1** pathways, are major accelerators of aging." |
| | AMPK, sirtuin 1 | "The other two nutrient sensors, **AMPK** and **sirtuins**, act in the opposite direction to IIS and mTOR, meaning that they signal nutrient scarcity and catabolism instead of nutrient abundance and anabolism. Accordingly, their up-regulation favours healthy aging. Moreover, **SIRT1** and **AMPK** can engage in a positive feedback loop, thus connecting both sensors of low-energy states into a unified response." |
| **Mitochondrial Dysfunction** | Mitochondrial dysfunction | "**Mitochondrial Dysfunction**" (Hallmark of Aging) |
| | Mitochondrial toxicity *(i)* | "Mitochondrial function becomes perturbed by aging-associated mtDNA mutations …". Another cause of perturbed mitochondrial function is toxin exposure, such as from alcohol, termed **mitochondrial toxicity**[4]. |
| | Reactive oxygen species (ROS) | "As chronological age advances, the levels of **ROS** increase in an attempt to maintain survival until they betray their original purpose and eventually aggravate, rather than alleviate, the age-associated damage." |

| | | |
|---|---|---|
| | Mitochondrial bioenergetics, mitochondrial biogenesis | "The reduced efficiency of **mitochondrial bioenergetics** with aging may result from multiple converging mechanisms including reduced **biogenesis of mitochondria**." |
| | Mitochondrial turnover, mitochondrial degradation | "The combination of increased damage and **reduced turnover in mitochondria**, due to lower biogenesis and **reduced clearance**, may contribute to the aging process." |
| | Electron transport chain (ETC), mitochondrial dynamics, Krebs cycle *(i)* | "Mitochondrial function becomes perturbed by aging-associated mtDNA mutations, reduced mitochondriogenesis, destabilization of the **electron transport chain (ETC)** complexes, altered **mitochondrial dynamics** or defective quality control by mitophagy." The **Krebs cycle** produces electron carriers that pass electrons to the electron transport chain (ETC). |
| **Cellular Senescence** | Cellular senescence | "**Cellular Senescence**" (Hallmark of Aging) |
| | Senescence markers | "The accumulation of senescent cells in aged tissues has been often inferred using **surrogate markers**, such as DNA damage. Some studies have directly used senescence-associated β-galactosidase (SABG) to identify senescence in tissues." |
| | Senescence-associated secretory phenotype (SASP) | "Senescent cells manifest dramatic alterations in their secretome, which … is referred to as the '**senescence-associated secretory phenotype'**." |
| | Immunosenescence (*i*) | Immunosenescence describes age-related alterations in the function of the immune system[5] and is partly explained by cellular senescence[6]. |
| **Stem Cell Exhaustion** | Stem cell exhaustion | "**Stem Cell Exhaustion**" (Hallmark of Aging) |
| | Progenitor cell, stem cell | "Although deficient proliferation of **stem** and **progenitor cells** is obviously detrimental for the long-term maintenance of the organism." |
| | Stem cell self-renewal | "Studies on aged mice have revealed an overall decrease in cell cycle activity of hematopoietic **stem cells** (HSCs), with old HSCs undergoing **fewer cell divisions** than young HSCs." |
| **Altered Intercellular Comm.** | Altered intercellular communication | "**Altered Intercellular Communication**" (Hallmark of Aging) |
| | Endocrine signalling, neuronal signalling, hormone *(i),* neurotransmitters *(i)* | "Aging also involves changes at the level of intercellular communication, be it **endocrine**, **neuroendocrine** or **neuronal**." In endocrine signalling, the signalling molecules are **hormones.** In neuronal signalling, the signalling molecules **neurotransmitters**. |
| | Inflammaging | "A prominent aging-associated alteration in intercellular communication is '**inflammaging'**." |
| | Inflammatory signalling | "Inflammaging may result from multiple causes such as the accumulation of **pro-inflammatory** tissue damage." |
| | Inflammation | "**Inflammation**" (Subheading) |

**Table S2. The original list of 207 ARDs, of which, 184 ARDs were included in the analysis**. The 23 ARDs that were excluded from further analysis are shown in italics including: (i) 4 ARDs that were not specific enough for scientific literature mining (*) and (ii) 19 ARDs with less than 250 associated publications (•). ARDs are categorised for clarity.

| ARD Category | List of ARDs |
|---|---|
| **Cancers** | Adrenal metastases |
| | Biliary cancer |
| | Bladder cancer |
| | Bone cancer |
| | Bone metastases |
| | Bowel cancer |
| | *Bowel metastases•* |
| | Brain cancer |
| | Brain metastases |
| | Breast cancer |
| | Kidney cancer |
| | Leukaemia |
| | Liver cancer |
| | Liver metastases |
| | Lung cancer |
| | Lung metastases |
| | Lymph node metastases |
| | Melanoma |
| | Mesothelioma |
| | Monoclonal gammopathy of undetermined significance (MGUS) |
| | Myelodysplasia |
| | Non-Hodgkin lymphoma (NHL) |
| | Oesophageal cancer |
| | Oropharyngeal cancer |
| | Ovarian cancer |
| | Pancreatic cancer |
| | Peritoneal metastases |
| | Plasma cell cancer |
| | Pleural metastases |
| | Polycythaemia vera |
| | *Primary malignancy (other)** |
| | Prostate cancer |
| | *Secondary malignancy (other)** |
| | Skin cancer |
| | Stomach cancer |
| | Thyroid cancer |
| | Uterine cancer |
| **Cardiovascular** | Abdominal aortic aneurysm (AAA) |
| | Atrial fibrillation (AF) |
| | Atrioventricular (AV) block (third degree) |
| | *Atrioventricular block (first degree)•* |
| | *Atrioventricular block (second degree)•* |
| | *Bifascicular block•* |
| | Cardiomyopathy |
| | Coronary heart disease (CHD) |
| | Dilated cardiomyopathy (DCM) |
| | Heart failure (CHF) |
| | Hypertension |
| | Hypertrophic cardiomyopathy (HCM) |
| | Intracerebral haemorrhage |

| | |
|---|---|
| | Ischaemic stroke<br>Left bundle branch block (LBBB)<br>*Multiple valve disorder•*<br>Myocardial infarction (MI)<br>Non-rheumatic aortic valve disorder<br>Non-rheumatic mitral valve disorder<br>Pericardial effusion<br>Peripheral arterial disease (PAD)<br>Primary pulmonary hypertension<br>Pulmonary embolism<br>Raynaud's disease<br>Rheumatic valve disorder<br>Right bundle branch block (RBBB)<br>*Secondary pulmonary hypertension•*<br>Sick sinus syndrome<br>Stable angina<br>Stroke<br>Subarachnoid haemorrhage (SAH)<br>Subdural haematoma<br>Supraventricular tachycardia (SVT)<br>Transient ischaemic attack (TIA)<br>*Trifascicular block•*<br>Unstable angina<br>Venous thromboembolism (VTE)<br>Ventricular tachycardia (VT) |
| **Benign Neoplasm** | *Benign brain neoplasms•*<br>*Benign colonic neoplasms•*<br>*Benign stomach neoplasms•* |
| **Digestive** | Abdominal hernia<br>*Angiodysplasia of colon•*<br>Anorectal prolapse<br>Autoimmune liver disease<br>Barrett's oesophagus<br>Cholangitis<br>Cholecystitis<br>Cholelithiasis<br>Cirrhosis<br>Diaphragmatic hernia<br>Diverticular disease<br>Fatty liver<br>Gastritis<br>Gastro-oesophageal reflux disease (GORD)<br>Liver failure<br>*Oesophageal ulcer•*<br>Oesophageal varices<br>Pancreatitis<br>Peptic ulcer<br>Peritonitis<br>Portal hypertension<br>Volvulus |
| **Endocrine** | Diabetes Mellitus (DM)<br>High (total) cholesterol<br>High triglycerides<br>Hyperparathyroidism |

| | |
|---|---|
| | Low high density lipoprotein cholesterol (low HDL)<br>Raised low density lipoprotein cholesterol (raised LDL)<br>Syndrome of inappropriate anti-diuretic hormone (SIADH)<br>Thyroid disease<br>Type 2 diabetes mellitus (T2DM) |
| **Ear** | Deafness<br>Meniere's disease<br>Tinnitus |
| **Eye** | Anterior uveitis<br>Blindness<br>Cataract<br>Diabetic eye disease<br>Glaucoma<br>Keratitis<br>Macular degeneration (AMD)<br>Ptosis<br>Retinal detachment<br>Retinal vascular occlusion |
| **Genitourinary** | Acute kidney injury (AKI)<br>Benign prostatic hyperplasia (BPH)<br>*Chronic cystitis•*<br>Chronic kidney disease (CKD)<br>End stage renal disease (ESRD)<br>Glomerulonephritis<br>Hydrocele<br>Neuropathic bladder<br>Obstructive and reflux uropathy<br>Tubulo-interstitial nephropathy<br>Urinary incontinence<br>Uterovaginal prolapse |
| **Haematological/ Immunological** | Agranulocytosis<br>Anaemia<br>Aplastic anaemia<br>*Folate deficiency anaemia•*<br>Haemolytic anaemia<br>Hypersplenism<br>*Hyposplenism•*<br>Immunodeficiency<br>Iron deficiency anaemia<br>Primary thrombocytopaenia<br>*Secondary polycythaemia•*<br>*Secondary thrombocytopaenia•*<br>*Vitamin B12 deficiency anaemia•* |
| **Infections** | Bacterial infection (ID)<br>Bone ID<br>Eye infection<br>Fungal infection<br>Gastroenteritis<br>Genitourinary infection (male)<br>Heart infection<br>*Infection of other organs**<br>*Infection with other organisms**<br>Lower respiratory tract infection (LRTI)<br>Nervous system infection |

| | |
|---|---|
| | Parasitic infection<br>Rheumatic fever<br>Sepsis<br>Skin infection<br>Urinary tract infection (UTI)<br>Viral infection |
| **Musculoskeletal** | Carpal tunnel syndrome<br>Collapsed vertebra<br>Fibromatosis<br>Giant cell arteritis<br>Gout<br>Hip fracture<br>Osteoarthritis<br>Osteoporosis<br>Polymyalgia rheumatica<br>Rheumatoid arthritis (RA)<br>Scleroderma<br>Scoliosis<br>Sjögren's Syndrome<br>Spinal stenosis<br>Spondylolisthesis<br>Spondylosis<br>Wrist fracture |
| **Neurological** | Autonomic neuropathy<br>Bell's palsy<br>Diabetic neuropathy<br>Epilepsy<br>Essential tremor<br>Motor neurone disease (MND)<br>Myasthenia gravis<br>Parkinson's disease<br>Peripheral neuropathy<br>Trigeminal neuralgia |
| **Psychiatric** | Delirium<br>Dementia |
| **Respiratory** | Asbestosis<br>Aspiration pneumonitis<br>Bronchiectasis<br>Chronic obstructive pulmonary disease (COPD)<br>Pleural effusion<br>*Pleural plaque*•<br>Pneumothorax<br>Pulmonary collapse<br>Pulmonary fibrosis<br>Respiratory failure |
| **Skin** | Actinic keratosis<br>Dermatitis<br>Lichen planus<br>Seborrheic dermatitis |

**Table S3. Examples of criteria for defining a sentence as a "confirmed association" between an aging hallmark and an ARD.** For example, if a sentence mentions that an ARD is: (i) caused or partially caused by, (ii) associated with, (iii) exacerbated by, or (iv) results in one of the criteria in the table below then a sentence is defined as "*confirmed association*" between an aging hallmark and ARD.

| Aging Hallmark | Criteria |
|---|---|
| Genomic Instability | • DNA damage or injury (e.g. DNA breaks)<br>• mitochondrial DNA damage or injury<br>• single or multiple somatic mutations<br>• genomic instability<br>• genetic alteration(s) |
| Telomere Attrition | • short or shorter leukocyte telomere length<br>• short or shorter telomere length<br>• telomere attrition, dysfunction or erosion<br>• telomere disorder |
| Epigenetic Alterations | • epigenetic alterations<br>• changes in DNA methylation<br>• changes in histone acetylation |
| Loss of Proteostasis | • proteasome dysfunction, dysregulation, failure or impairment<br>• aggregated or misfolded proteins<br>• induction of, or increased endoplasmic reticulum (ER) stress<br>• activation of, or aberrant unfolded protein response (UPR)<br>• inadequate, dysregulated or defective autophagy<br>• impaired proteostasis<br>• impaired chaperone activity<br>• reduced proteolytic activity |
| Deregulated Nutrient Sensing | • increased insulin resistance<br>• dyslipidaemia<br>• decreased 5' AMP-activated protein kinase (AMPK) activity<br>• decreased sirtuin 1 activity |
| Mitochondrial Dysfunction | • generation or presence of reactive oxygen species (ROS) or active oxygen<br>• mitochondrial dysfunction or degeneration<br>• altered mitochondrial dynamics or bioenergetics<br>• mitochondrial damage<br>• impaired mitochondrial turnover including mitochondrial biogenesis or mitophagy |
| Cellular Senescence | • accelerated, early or enhanced cellular senescence, replicative senescence or immunosenescence<br>• activation of cellular senescence or cellular aging<br>• association with the senescence-associated secretory phenotype (SASP)<br>• increased senescent cells or expression of senescence markers |
| Stem Cell Exhaustion | • reduced number, impaired proliferative capacity or increased destruction of stem cells or progenitor cells<br>• impaired function, mobilization or exhaustion of stem cells or progenitor cells<br>• aging or senescence of stem cells or progenitor cells<br>• decreased number of circulating stem cells or progenitor cells<br>• decreased stem cell or progenitor cell differentiation<br>• stem cell disorder |
| Altered Intercellular Communication | • increased inflammation<br>• decreased levels of specific hormones, e.g. oestradiol, testosterone<br>• decreased synaptic transmission<br>• increased levels of specific hormones, e.g. parathyroid hormone |

**Table S4. Examples of sentences correctly reporting that an aging hallmark (yellow) has a role in the development or disordered physiology of an ARD (grey)**. Sentences such as these were labelled as a "confirmed association" between an aging hallmark and an ARD on manual curation. Those aging hallmark-ARD combinations with insufficient evidence were set to zero. Table S9 provides a list of abbreviations.

| Aging Hallmark | ARD (MeSH ID) | PubMed ID | Sentence |
|---|---|---|---|
| GI (DNA damage) | Motor neurone disease (D016472, D000690, D010244) | 18344116 | "Increased mitochondrial oxidative damage and oxidative DNA damage contributes to the neurodegenerative process in sporadic amyotrophic lateral sclerosis."[7] |
| TA (Decreased telomere length) | Aplastic anaemia (D000741, D029502, D005199, D029503) | 22687638 | "Telomere shortening, a well-known marker of aging and cellular stress, occurs under several conditions in the hematopoietic compartment, including aplastic anemia and following iatrogenic noxae."[8] |
| EA (Epigenetic alterations) | Barrett's oesophagus (D001471) | 29046735 | "Identification of a key role of widespread epigenetic drift in Barretts oesophagus and oesophageal adenocarcinoma."[9] |
| LOP (Protein aggregation) | Cataract (D058442, D002386) | 23179275 | "Cataract, the loss of transparency of eye lens, is a disease of protein aggregation."[10] |
| DNS (Insulin resistance) | Type 2 diabetes mellitus (D003924) | 15579187 | "Insulin resistance is the central defect in the development of type 2 diabetes, preceding its onset by 10-20 years."[11] |
| MD (Reactive oxygen species) | Dementia (D003704, D000544, D015140, D015161) | 20933077 | "Mitochondrial oxidative stress induced by reactive oxygen species (ROS) has been strongly associated with the pathogenesis of neurodegenerative disorders, including Alzheimer's disease (AD)."[12] |
| CS (Cellular senescence) | Diabetic retinopathy (D003930) | 25162034 | "Accumulating evidence has shown that diabetes accelerates aging and endothelial cell senescence is involved in the pathogenesis of diabetic vascular complications, including diabetic retinopathy."[13] |
| SCE (Progenitor cell) | Osteoarthritis (D010003, D015207, D020370, D055013) | 21815581 | "The lower chondrogenic and spontaneous osteogenic differentiation of mesenchymal progenitor cells derived from elderly patients may be associated with the development of primary osteoarthritis."[14] |
| AIC (Inflammation) | Myocardial infarction (D009203, D056988, D056989, D000072658, D0000726657) | 10347344 | "Inflammation plays a critical role in acute myocardial infarction (AMI) and tumor necrosis factor alpha (TNF-alpha) is a potent inflammatory trigger."[15] |

**Table S5**. **Summary of literature on each aging hallmark in the human aging corpus**. The number of (a) abstracts and (b) sentences mentioning each aging hallmark. (c) The number of sentences also mentioning ARDs per aging hallmark (i.e. co-mentioning the aging hallmark and any ARD).

| Aging hallmark | (a) Number of abstracts | (b) Number of sentences | (c) Number of sentences with co-mentions |
|---|---|---|---|
| Genomic Instability | 7,313 | 12,975 | 2,166 |
| Telomere Attrition | 1,823 | 6,669 | 864 |
| Epigenetic Alterations | 10,751 | 34,403 | 8,333 |
| Loss of Proteostasis | 5,748 | 13,915 | 1,694 |
| Mitochondrial Dysfunction | 5,737 | 11,785 | 1,257 |
| Deregulated Nutrient Sensing | 15,753 | 34,227 | 12,682 |
| Cellular Senescence | 4,896 | 10,173 | 511 |
| Stem Cell Exhaustion | 7,466 | 15,068 | 2,344 |
| Altered Intercellular Comm. | 77,783 | 133,733 | 25,958 |

**Table S6**. **Features of the ARD multimorbidity networks.** The table provides a summary of the 4 ARD multimorbidity networks including the number of nodes and average network density. We derived subnetworks of the top 30 ARDs for each aging hallmark from these multimorbidity networks.

| Age category | Number of ARD nodes | Average network density |
|---|---|---|
| 50 – 59 years | 184 | 0.0999 |
| 60 – 69 years | 184 | 0.0976 |
| 70 – 79 years | 184 | 0.0836 |
| ≥80 years | 184 | 0.0684 |

**Table S7. Network density of subnetworks of the top 30 ranking ARD nodes after network propagation for age categories 50-59 years, 60-69 years, 70-79 years and ≥80 years.** The number of times the network density from permutations (n = 20,000) was greater than or equal to the true network density for that aging hallmark was used to calculate the p-value. The p-value was corrected for multiple testing across the 4 age categories per aging hallmark using the Benjamini-Hochberg procedure (*p < 0.05, ** p <0.01, *** p <0.001, ****p<0.0001).

| Aging Hallmark | ARD Network Density | | | |
|---|---|---|---|---|
| | 50-59 years | 60-69 years | 70-79 years | ≥80 years |
| Genomic Instability | 0.1425 | 0.1402 | 0.1310 | 0.1241 |
| Telomere Attrition | 0.1195 | 0.1356 | 0.1195 | 0.1034 |
| Epigenetic Alterations | 0.2000** | 0.1885** | 0.2092*** | 0.1885*** |
| Loss of Proteostasis | 0.0897 | 0.0805 | 0.0759 | 0.0621 |
| Deregulated Nutrient Sensing | 0.3448**** | 0.3632**** | 0.3448**** | 0.3034**** |
| Mitochondrial Dysfunction | 0.1931**** | 0.1931*** | 0.1425*** | 0.1057** |
| Cellular Senescence | 0.1563 | 0.1609 | 0.1379 | 0.1034 |
| Stem Cell Exhaustion | 0.2322**** | 0.2322*** | 0.2161*** | 0.1931*** |
| Altered Intercellular Comm. | 0.2207*** | 0.2161*** | 0.1977*** | 0.1540** |

**Table S8. (a) Inclusion and (b) exclusion criteria were applied when defining the "human aging corpus" to increase relevance of the selected abstracts.**

| a) Inclusion Criteria | Rationale |
|---|---|
| Journal articles | Articles were required to be indexed with the "journal article" publication type. |
| Publication date | Journal articles published between the date of the landmark paper on lifespan extension, "*A C. elegans mutant lives twice as long as wildtype*", on 2nd December 1993 to 31st December 2017 were included. |
| English language | Abstracts were required to be in English. |
| Humans | We confined our search to the literature on humans because certain mechanisms of aging are only found in particular evolutionary lineages[16]. Therefore, articles were required to be indexed with the humans Medical Subject Heading (MeSH) term, as certain mechanisms of aging are private. |
| Aging synonyms | Articles were required to be associated with one of 43 aging terms. |
| **b) Exclusion Criteria** | **Rationale** |
| Specific publication types | Reviews, introductory journal articles, retractions of publication, letters to the editor and comments were excluded in order to retrieve high quality original research rather than data from secondary sources. We excluded review articles to avoid re-using content within the scientific literature. |
| Germline mutation | Abstracts mentioning, or indexed with, 'germline mutations' or related terms were excluded. Germline mutations can be a consequence of the maternal or paternal aging process. Thus, germline mutations were excluded to maximize retrieval of articles focused on the effects of the individual's own aging process. |
| Telomerase activation and inhibition | Abstracts mentioning 'telomerase activation', 'telomerase inhibition' or related terms were excluded. Telomerase activation is a consequence of cancer progression, as opposed to aging, and leads to telomere lengthening. Therefore, telomerase activation and inhibition were excluded to reduce selection of irrelevant abstracts under the telomere attrition aging hallmark. |
| Adverse drug effects | Abstracts reporting adverse drug effects, such as DNA damage from chemotherapy for cancer, were excluded to reduce selection of irrelevant abstracts. |
| Hormone preparations and anti-inflammatory drugs | Abstracts mentioning 'hormone preparations', 'anti-inflammatory drugs' or related terms, were excluded to reduce selection of irrelevant abstracts under the altered intercellular communication aging hallmark. |
| Stem cell transplant | Abstracts mentioning 'stem cell transplant', or related terms, were excluded to reduce selection of irrelevant abstracts under the stem cell exhaustion aging hallmark. |
| Induced Pluripotent stem cells (iPSC) | Abstracts mentioning, or indexed with, induced pluripotent stem cells (iPSC) or related terms were excluded. iPSC are used experimentally to treat ARDs, which can result in incorrect selection of abstracts under the stem cell exhaustion aging hallmark. |
| Neoplastic stem cells | Abstracts mentioning, or indexed with, 'neoplastic stem cells' or related terms were excluded. According to the cancer stem-cell hypothesis, neoplastic stem cells are responsible for cancer tumour growth. This contrasts with stem cell exhaustion, where loss of stem cells leads to ARD development. Therefore, neoplastic stem cells were excluded to avoid incorrect selection of abstracts under the stem cell exhaustion aging hallmark. |
| GWAS catalog | We excluded PMIDs that were linked to studies included from the GWAS catalog. Thus, the genetic approach to verifying aging hallmark-ARD associations was independent of the literature-based method. |

**Table S9. Table of abbreviations.** Abbreviations are sorted in alphabetical order for (a) aging hallmarks, (b) age-related diseases and (c) genes & proteins.

| Abbreviation | Term |
|---|---|
| (a) Aging hallmark abbreviations | |
| AH | Aging hallmark |
| AIC/ Altered intercellular comm. | Altered intercellular communication |
| AMPK | 5’ AMP-activated protein kinase |
| CS | Cellular senescence |
| DNA | Deoxyribonucleic acid |
| DNS | Deregulated nutrient sensing |
| ds-DNA | Double stranded DNA |
| EA | Epigenetic alterations |
| ER | Endoplasmic reticulum |
| GI | Genomic instability |
| IIS pathway | Insulin/insulin like growth factor (IGF)-1 signalling pathway |
| LOP | Loss of proteostasis |
| LTL | Leukocyte telomere length |
| MD | Mitochondrial dysfunction |
| mtDNA | Mitochondrial DNA |
| mTORC1 | mammalian Target Of Rapamycin Complex 1 |
| RNA | Ribonucleic acid |
| ROS | Reactive oxygen species |
| SASP | Senescence associated secretory phenotype |
| SCE | Stem cell exhaustion |
| ss-DNA | Single stranded DNA |
| TA | Telomere attrition |
| TL | Telomere length |
| UPR | Unfolded protein response |
| (b) Age-related disease abbreviations | |
| AAA | Abdominal aortic aneurysm |
| AF | Atrial fibrillation |
| AKI | Acute kidney injury |
| AMD | Age-related macular degeneration |
| ARD | Age-related disease |
| Autoimmune liver Dz. | Autoimmune liver disease |
| Bacterial ID | Infectious disease, bacterial |
| Barrett’s | Barrett’s oesophagus |
| Bone ID | Infectious disease of the bone |
| BPH | Benign prostatic hyperplasia |
| CHD | Coronary heart disease |
| CHF | Congestive heart failure |
| CKD | Chronic kidney disease |
| COPD | Chronic obstructive pulmonary disease |
| DCM | Dilated cardiomyopathy |
| Diabetic Eye Dz. | Diabetic eye disease/ diabetic retinopathy |
| DM | Diabetes mellitus |
| ESRD | End stage renal disease |
| GORD | Gastro-oesophageal reflux disease |
| HCM | Hypertrophic cardiomyopathy |
| Low HDL | Low high-density lipoprotein cholesterol |
| LRTI | Lower respiratory tract infection |
| Mets. | Metastases |

| | |
|---|---|
| MGUS | Monoclonal gammopathy of undetermined significance |
| MI | Myocardial infarction |
| MND | Motor neurone disease |
| NHL | Non-Hodgkin's lymphoma |
| PAD | Peripheral arterial disease |
| Parkinson's | Parkinson's disease |
| Pri. thrombocytopaenia | Primary thrombocytopaenia |
| RA | Rheumatoid arthritis |
| Raised LDL | Raised low density lipoprotein (LDL) cholesterol |
| Retinal vascular occ. | Retinal vascular occlusion |
| SAH | Subarachnoid haemorrhage |
| SIADH | Syndrome of inappropriate anti-diuretic hormone (ADH) |
| Sjogren's | Sjogren's syndrome |
| T2DM | Type 2 diabetes mellitus |
| TIA | Transient ischaemic attack |
| UTI | Urinary tract infection |
| VTE | Venous thromboembolism |
| (c) Gene & protein abbreviations | |
| ABCA7 | ATP binding cassette subfamily A member 7 |
| AGER | Advanced glycosylation end-product specific receptor |
| ANGPT1 | Angiopoietin 1 |
| ATAD5 | ATPase family AAA domain containing 5 |
| BAG6 | BAG cochaperone 6 |
| BCAR1 | BCAR1 scaffold protein, Cas family member |
| BCL3 | BCL3 transcription coactivator |
| BRAF | B-Raf proto-oncogene, serine/threonine kinase |
| BRCA2 | BRCA2 DNA repair associated |
| BTN3A1 | Butyrophilin subfamily 3 member A1 |
| CAMK2(B/D/G) | Calcium/calmodulin dependent protein kinase II (beta/ delta/ gamma) |
| CD(28/36/247/226) | CD28/ CD36/ CD247/ CD226 molecule |
| CDKN1A | Cyclin dependent kinase inhibitor 1A |
| CFLAR | CASP8 and FADD like apoptosis regulator |
| CHEK2 | Checkpoint kinase 2 |
| CHUK | Component of inhibitor of nuclear factor kappa B kinase complex |
| CTNNA3 | Catenin alpha 3 |
| DENND1B | DENN domain containing 1B |
| DSTYK | Dual serine/threonine and tyrosine protein kinase |
| ERK1/2 | Mitogen-activated protein kinase 3/ 1 |
| FBXW7 | F-box and WD repeat domain containing 7 |
| FFAR4 | Free fatty acid receptor 4 |
| FGA | Fibrinogen alpha chain |
| FGB | Fibrinogen beta chain |
| FGF10 | Fibroblast growth factor 10 |
| FGFR2 | Fibroblast growth factor receptor 2 |
| FGFR3 | Fibroblast growth factor receptor 3 |
| FGFR4 | Fibroblast growth factor receptor 4 |
| GAREM1 | GRB2 associated regulator of MAPK1 subtype 1 |
| GAS6 | Growth arrest specific 6 |
| GATA3 | GATA binding protein 3 |
| GO | Gene Ontology |

| | |
|---|---|
| GPNMB | Glycoprotein Nmb |
| GPR183 | G protein-coupled receptor 183 |
| GSEA | Gene Set Enrichment Analysis |
| GWAS | Genome Wide Association Study |
| HAND2 | Heart and neural crest derivatives expressed 2 |
| HLA-B | Major histocompatibility complex, class I, B |
| HLA-DPA1 | Major histocompatibility complex, class II, DP alpha 1 |
| HLA-DQB1 | Major histocompatibility complex, class II, DQ beta 1 |
| HLA-DR(B1/A/B5) | Major histocompatibility complex, class II, DQ beta 1/ alpha/ beta 5 |
| HMGCR | 3-hydroxy-3-methylglutaryl-CoA reductase |
| IFN-$\gamma$ | Interferon gamma |
| IFNGR2 | Interferon gamma receptor 2 |
| INPP5D | Inositol polyphosphate-5-phosphatase D |
| IRF(4/5/8) | Interferon regulatory factor 4/5/8 |
| JAK2 | Janus kinase 2 |
| KCNN4 | Potassium calcium-activated channel subfamily N member 4 |
| LINGO1 | Leucine rich repeat and Ig domain containing 1 |
| LRRTM3 | Leucine rich repeat transmembrane neuronal 3 |
| NAT2 | N-acetyltransferase 2 |
| NECAB2 | N-terminal EF-hand calcium binding protein 2 |
| NECTIN2 | Nectin cell adhesion molecule 2 |
| NELFE | Negative elongation factor complex member E |
| NFKB1 | Nuclear factor kappa B subunit 1 |
| NOD2 | Nucleotide binding oligomerization domain containing 2 |
| NOTCH2 | Notch receptor 2 |
| OASL | 2'-5'-oligoadenylate synthetase like |
| OPRM1 | Opioid receptor mu 1 |
| PAK2 | p21 (RAC1) activated kinase 2 |
| PDE4D | Phosphodiesterase 4D |
| PDGFC | Platelet derived growth factor C |
| PHLDA3 | Pleckstrin homology like domain family A member 3 |
| PLC(G1/G2) | Phospholipase C gamma 1/ 2 |
| PRKD2 | Protein kinase D2 |
| PTK2B | Protein tyrosine kinase 2 beta |
| PTPN11/22 | Protein tyrosine phosphatase non-receptor type 11/22 |
| RAB29 | RAB29, member RAS oncogene family |
| RAF1 | Raf-1 proto-oncogene, serine/threonine kinase |
| RAP1A | RAP1A, member of RAS oncogene family |
| RASGRP1 | RAS guanyl releasing protein 1 |
| SCIMP | SLP adaptor and CSK interacting membrane protein |
| SKAP1 | src kinase associated phosphoprotein 1 |
| TAB2 | TGF-beta activated kinase 1 (MAP3K7) binding protein 2 |
| TGFB1 | Transforming growth factor beta 1 |
| TNFRSF11A | TNF receptor superfamily member 11a |
| TP53 or p53 | Tumor protein p53 |
| TP63 | Tumor protein p63 |
| TREM2 | Triggering receptor expressed on myeloid cells 2 |
| TRIM(5/8/26/31) | Tripartite motif containing 5/8/26/31 |
| VEGFA | Vascular endothelial growth factor A |
